# *In vivo* single-molecule imaging of RecB reveals efficient repair of DNA damage in Escherichia coli

**DOI:** 10.1101/2023.12.22.573010

**Authors:** Alessia Lepore, Daniel Thédié, Lorna McLaren, Benura Azeroglu, Oliver J. Pambos, Achillefs N. Kapanidis, Meriem El Karoui

## Abstract

Efficient DNA repair is crucial for maintaining genome integrity and ensuring cell survival. In *Escherichia coli*, RecBCD plays a crucial role in processing DNA ends following a DNA double-strand break (DSB) to initiate repair by homologous recombination. While RecBCD has been extensively studied *in vitro*, less is known about how it contributes to rapid and efficient repair in living bacteria. Here, we perform single-molecule microscopy to investigate DNA repair in real-time in *E. coli*. We quantify RecB single-molecule mobility and monitor the induction of the DNA damage response (SOS response) in individual cells. We show that RecB binding to broken DNA ends leads to efficient repair without SOS induction. In contrast, in a RecB mutant with modified activities leading to the activation of an alternative repair pathway, repair is less efficient and leads to high SOS induction. Our findings reveal how subtle alterations in RecB activity profoundly impact the efficiency of DNA repair in *E. coli*.

## 1 INTRODUCTION

DNA repair is a fundamental mechanism that ensures chromosome maintenance and cell survival after DNA damage [1]. Among the different kinds of DNA lesions, DNA Double Strand Breaks (DSBs) are one of the most threatening to genome stability. Unrepaired DSBs can lead to cell death, while incomplete or faulty repair can induce mutagenesis and genome rearrangement [2]. DSBs can be caused by endogenous or exogenous causes such as the collapse or stalling of replication forks, oxygen radicals, ionizing radiation and DNA-damaging agents [3]. DNA replication is the main physiological source of DSB. In *Escherichia coli*, 18% of cells per generation experience spontaneous replication fork breakage [4]. Quinolone antibiotics, which target DNA-topoisomerase, disrupt DNA replication to induce DSBs, ultimately leading to bacterial cell death. This class of antibiotic, binding to the topoisomerase-DNA complex, interferes with changes in the DNA supercoiling and causes the arrest of the replication machinery. Consequently, to remove the block on the replication fork, DSBs are formed [5]. However, DSBs can be repaired through homologous recombination, in which the missing information is copied from another intact, identical chromosome [6].

In *E. coli*, the initial phase of the repair pathway involves the heterotrimer complex RecBCD [3]. This complex plays a crucial role in repairing DSBs by binding to DNA ends and processing them for subsequent homologous recombination. RecBCD is expressed at very low levels in cells [7] and the regulation of its expression levels is crucial for the cell’s DNA repair capability [8]. Although RecBCD expression is not regulated by DNA damaging events, both deletion and over-expression of RecBCD strongly affect DNA repair, cell viability and homologous recombination [9, 10, 11]. After it locates the DNA ends, RecBCD utilizes its two helicase motors with distinct polarities, namely RecB with a 3’ *→* 5’ direction and RecD with a 5’ *→* 3’ direction, to translocate along both DNA strands. During this translocation process, RecB’s nuclease activity actively degrades both DNA strands until it encounters a specific octameric DNA sequence known as *χ*-site (5’-GCTGGTGG-3’). The recognition of the *χ*-site triggers a modulation in RecBCD’s biochemical activities, leading to a drastic reduction in RecB’s nuclease activity at the 3’ single-stranded DNA region. This alteration facilitates the loading of the RecA protein onto the 3’ single-stranded DNA tail by RecBCD, forming a RecA-ssDNA filament. The RecA filament then catalyses homology search and strand invasion, facilitating replication restart.

RecA binding to single-stranded DNA also triggers LexA autoproteolysis, which activates the SOS regulon genes, allowing *E. coli* to respond to and repair DNA damage [12]. The SOS regulon comprises approximately 40 genes. Among the genes regulated by LexA are DNA repair genes such as RecA, and inhibitors of cell division e.g. SulA [13, 12]. Inhibition of cell division by SulA results in bacterial cells appearing as elongated with an increased cell area in comparison to cells without DNA damage [14].

*In vitro* studies have not only demonstrated the crucial role of RecBCD activity in recognizing and processing damaged DNA ends but have also highlighted its significance in ensuring the successful formation of the RecA ssDNA filament [15, 16, 17, 18, 19, 20]. The study of RecBCD crystal structure bound to a DNA hairpin allowed an understanding of how RecBCD interacts with DNA clarifying the fate of the 3’ ssDNA after the RecC domain recognises the *χ*-site. While RecBCD keeps degrading the 5’ side and translocating on the DNA, RecA is recruited on the forming ssDNA loop [15]. Interestingly, RecA recruitment has been associated with the presence of the RecB nuclease domain. RecBCD mutants in which the RecB nuclease function is inactivated fail to recruit RecA to the 3’ ssDNA [21, 22, 23]. In particular, the *recBD1080A* mutant (known as *recB1080*) contains a single point mutation that inactivates the nuclease domain. *In vitro* data shows that RecB1080 is a functional helicase that unwinds DNA without degrading the 3’ ssDNA. However, it has been observed that, while it still recognizes *χ* sites, it does not promote RecA loading onto the ssDNA.

The precise mechanism by which RecBCD disengages from the DNA remains to be fully elucidated. *In vitro* observations have led to the formulation of a model [24], suggesting that the RecBCD dissociation process is initiated after recognition of the *χ*-site. According to this hypothesis, post *χ*-site recognition, RecBCD continues to unwind the DNA beyond the *χ*-site and then the subunits disengage from the DNA. In this model, the possible impact of the RecA filament formation on RecBCD-DNA dissociation is not taken into account. However, considering the role of the RecB nuclease domain in recruiting RecA to the ssDNA and the intricate topological shape of the RecBCD-DNA-RecA complex, it may play a role in RecBCD disengagement from the DNA.

While *in vitro* studies have laid the foundations of the mechanisms of the repair and the enzyme’s activities, *in vivo* observations gain a deeper understanding of how DSB repair happens in the complex environment of the live cell. The homologous recombination event following DSB repair has been observed in live *E. coli*. Employing various methods to cause DNA DSBs, RecA filament formation and activity have been monitored using different fluorescent fusions and labelling techniques [25, 26, 27, 28]. In a recent live-cell study, the observation of the disappearance of fluorescent loci placed on the DNA after DSB induction confirmed that, in *E. coli*, RecBCD exhibits high translocating speed on DNA (up to *∼*1.6 kb/s) and high degradation activity (*∼*100 kb) [29]. Although recent work explored RecB mobility after mitomycin C treatment [30], it is still unclear how RecBCD dynamics change in response to various levels of DNA damage. Understanding RecBCD dynamics at different levels of DNA damage is crucial to reveal how different amounts of DSBs are detected and processed *in vivo*.

To characterize RecBCD-mediated DSBs repair *in vivo*, we observed and quantified the mobility of single RecB molecules in live *E. coli* in real-time. RecB has a crucial role in functionality since it is the only subunit of the complex that acts as both nuclease and helicase, and as such, RecB is an excellent candidate to study RecBCD activities. We induced different levels of DNA damage using a fluoroquinolone antibiotic, ciprofloxacin, while monitoring the SOS response in the bacterial population. We observed how the RecB molecular mobility changes at different DNA damage levels. We identified three sub-populations of RecB molecules, each describing RecB mobility within bacterial cells. Furthermore, we determined that the fraction of RecB molecules involved in the repair process is proportional to the level of DNA damage. To explore the impact on DBS repair when RecBCD cannot promote RecA filament formation, we quantified the mobility and the SOS induction in the *recB1080* mutant. Our observations are consistent with a model based on previous *in vitro* observations [23], which suggests an alternative pathway for loading RecA onto single-stranded DNA when RecB-mediated RecA loading is impaired. This implies that *in vivo*, the alternative repair pathway operates on a longer time scale and with reduced efficiency compared to the repair process in the WT.

## 2 MATERIALS AND METHODS

### 2.1 Strains and plasmid construction

*E. coli* MG1655 strain and its derivatives were used in this study. The characteristics of all the strains and plasmids employed are described in Table 1. The construction of the strain carrying the RecB-HaloTag fusion (MEK65) is described in [7]. To build the strains containing the GFP expression reporter (MEK707 and MEK2324), *PsulA-mGFP* was cloned into a pOSIP plasmid [31] and integrated at a chromosomal locus into the genome by clone integration [31]. The construction of the pOSIP plasmid containing the fluorescent reporter (pSJR036) is described in [32]. After construction, the MEK707 and MEK2324 strains were checked by PCR amplification of the insertion (see Supp. Table 1). The *recB1080* mutation was introduced into the *recB-HaloTag* strain to create the *recB1080-HaloTag* strain (MEK716). This was achieved through plasmid-mediated gene replacement using a plasmid derived from pTOF24, pDL4174 [33]. The *recB1080* fragment was generated by PCR with the primers listed in Supplementary Table 1. A digestion site, HaeIII, was incorporated into the *recB1080* fragment to facilitate PCR verification of successful construction. Subsequently, the *recB1080* fragment was ligated into a pTOF24 backbone after the plasmid was digested at the PstI and SalI sites, resulting in pDL4174. After construction, MEK 716 was checked by restriction-digestion with HaeIII enzyme and PCR amplification.

The nucleotide sequences used are listed in Supplementary Table 1.

**Table 1:** List of strains and plasmids used in this work.

| Name | Characteristics | Reference |
| --- | --- | --- |
| **Bacterial Strain** |  |  |
| MG1655 | *F-lambda-rph-1* | Lab stock |
| MEK65 | MG1655,<br>*recB::halotag* | [7] |
| MEK445 | MG1655,<br>*HK022::psfiA-GFP* | [32] |
| MEK707 | MG1655,<br>*recB::halotag*<br>*HK022::psfiA-GFP* | this work |
| MEK716 | MG1655,<br>*recB1080::halotag* | this work |
| DL654 | MG1655,*recA::CmR* | [34] |
| MEK1326 | MG1655, $\Delta recB$ | [8] |
| MEK2324 | MG1655,<br>*recB1080::halotag*<br>*HK022::psfiA-GFP* | this work |
| **Plasmids** |  |  |
| pSJR036 | *PsulA-mGFP* in-<br>sertion by Clone<br>integration | [32] |
| pDL4174 | pTOF with *recB1080* | this work,<br>gift from<br>the Leach<br>Lab |

### 2.2 Antibiotic susceptibility tests

Antibiotic susceptibility tests were conducted by cultivating the relevant bacterial strains (MG1655, MEK65, MEK707, MEK716, MEK2324, MEK1326, DL654) overnight in LB media at 37^*°*^C. From the overnight cultures, serial dilutions were prepared with a dilution factor of 10^*−*5^ starting from OD_600_ = 1, and each strain was subsequently plated using a 42-pinner onto LB agar plates containing varying concentrations of ciprofloxacin (0, 4, 10, 14, 16, 20 ng/ml). These plates were then incubated overnight at 37^*°*^C (see Supp. Figure 8).

### 2.3 Culture conditions and Halo Labelling

For all microscopy-based experiments, cells were grown in M9 supplemented with 0.2% (w/v) of glucose, 2 mM MgSO_4_, 0.1 mM CaCl_2_, 1*×* MEM Essential and MEM Non-Essential Amino Acids (Gibco®). For single-molecule labelling, we used the labelling protocol presented in [7] without the chemical fixation step. In brief, bacterial cultures from frozen -80^*°*^C stocks were grown with shaking (150 rpm) in the culture medium overnight (14-16 hours) at 37^*°*^C. The overnight cultures were diluted (1:300) into 15 ml of medium and grown at 37^*°*^C to the mid-exponential phase (optical density OD_600_ = 0.2–0.3). A volume of cells equivalent to 1 mL at OD_600_ = 0.2 was centrifuged and re-suspended in 1ml fresh medium supplemented with JF549 (Janelia Fluor®HaloTag®Ligands, Promega) at a final concentration of 1 *µ*M. The culture was further incubated for 1 hour at 37^*°*^C with shaking. After the labelling step, each sample was centrifuged for 3 min at 8000 rpm and the pellet was resuspended in 0.5–1 ml of the M9-based medium (dye-free). This washing step was repeated 3–4 times. At each step, cells were transferred to a new tube to facilitate the removal of the dye. After the last washing step, 2–2.5 *µ*l of bacteria were added to an agar pad containing 2% agarose dissolved in M9 media.

To induce DNA damage, ciprofloxacin at the chosen concentration (4, 10 or 14 ng/ml) was added to the bacterial culture 150 min before microscopy. The same concentration was maintained in the agar pad during microscopy.

#### 2.3.1 SYTOX labelling

The bacterial nucleoid of the *recB-HaloTag* strain MEK65, which does not carry the GFP SOS reporter, was labelled using SYTOX green (Invitrogen) [35]. During the Halo labelling protocol, 500 nM of SYTOX green was added 40–45 min before the beginning of the washing step.

### 2.4 Microscopy set-up

Imaging was performed using an inverted microscope (Nikon Ti-E) equipped with an EMCCD Camera (iXion Ultra 897, Andor), a 100X TIRF Nikon objective (NA 1.49, oil immersion) and a 1.5X Nikon magnification lens (pixel size = 107 nm). Images were acquired via MetaMorph®(Molecular Devices; v7.8.13.0) in HILO (Highly Inclined Laminar Optical sheet) [36] configuration. The HILO configuration was established using the iLas® variable angle TIRF control window.

### 2.5 Real-time DNA repair imaging

Fluorescence excitation was performed using 561 nm and 488 nm lasers (Coherent OBIS) and detected via a dual-wavelength dichroic filter (488/561 nm) (TRF59904, Chroma). This configuration was used to stimulate the fluorescent emissions from RecBHaloTag-JF549 and the fluorescent signal emitted by *PsulA-GFP* reporter. Movies were acquired with continuous laser excitation at 561 nm at *∼* 15mW with an exposure time of 12 ms for a total acquisition time of 7 sec (600 frames). The camera’s electron-multiplying (EM) gain was set to 300, and the region of interest (ROI) was set to 256x256 pixels. A snapshot of the same ROI was acquired to image SOS induction by exciting the GFP signal with a 488 nm laser at *∼* 6 mW for 80 ms (camera EM gain 50). For each ROI, bright-field z-stacks of 16 images were acquired around the focus (total distance 3 *µ*m, each step of 0.2 *µ*m). Each bright-field image was acquired with 30 ms of exposure time and an EM camera gain of 4. When nucleoid images were acquired instead of the SOS induction signal, the SYTOX green GFP signal was excited with the 488 nm laser at *∼* 6 mW for 30 ms (camera EM gain 4). Each sample was imaged for a maximum time of 40-45 min. All the acquisitions were performed at 37^*°*^C in an Okolab microscope cage incubator equipped with dark panels.

#### 2.5.1 Cell segmentation

Cell segmentation was performed from bright-field images in BACMMAN [37], an ImageJ plug-in for high-throughput image analysis and manual curation. Bright-field images were first imported into BACMMAN as a “Dataset”. The “pre-processing” step was then applied, which consisted of a single step that cropped the 16-image bright-field z-stack to 5 images on one side of the focus, as required by our cell segmentation algorithm. In the next step of the pipeline (the “processing” step), cells were segmented using Talissman, a U-net-based segmentation algorithm (https://github.com/jeanollion/TaLiSSman). In brief, the U-net model predicts an Euclidean distance map, where the value of each pixel is its predicted distance to the nearest background pixel. A watershed algorithm is then applied to retrieve cell contours. This approach allowed us to accurately segment cells from bright-field images, including when they formed tight clusters. Following segmentation, post-filters were applied to dilate the segmented regions slightly (to make sure we would not miss any fluorescent spots located near the edge of the cell during single-particle tracking) and to remove any cells that were in contact with the edge of the image and might therefore be cropped. The resulting segmentation masks were finally exported in hdf5 format and imported to MATLAB to resolve cells during single-particle tracking.

#### 2.5.2 Nucleoid detection

SYTOX green fluorescence images were analysed in BACMMAN. First, a deep-learning-based denoising algorithm [38] was applied. Individual nucleoids were then segmented using a watershed algorithm on the maximum eigen-values of the Hessian transform of the image. This approach allows precise segmentation of large spot-like objects with variable shapes. The segmented regions were exported in hdf5 format for further processing.

#### 2.5.3 SOS induction signal detection and quantification

After performing bacterial cell segmentation, we computed the average fluorescent signal for each segmented cell in BACMMAN. The local fluorescent background was subtracted from each pixel of the image during the “preprocessing” step using the ImageJ background subtraction method (Class: BackgroundSubstracted).

### 2.6 Comparison of SOS induction in the WT and the RecBHaloTag

All the data were acquired and analysed as described above, except for the data shown in Supplementary Figures 1C and D, which were acquired and analysed as described below. Bacterial cells were grown in the same media described previously. Following overnight incubation, bacterial cultures were diluted (1:1000) into 15 ml of the medium and grown at 37^*°*^C until the OD_600_ reached 0.4. Ciprofloxacin was added when necessary at a concentration of 10 ng/ml, and the incubation continued for a total of 150 minutes. Samples were imaged on M9-based agar pads consisting of 2% agarose.

**Figure 1.**
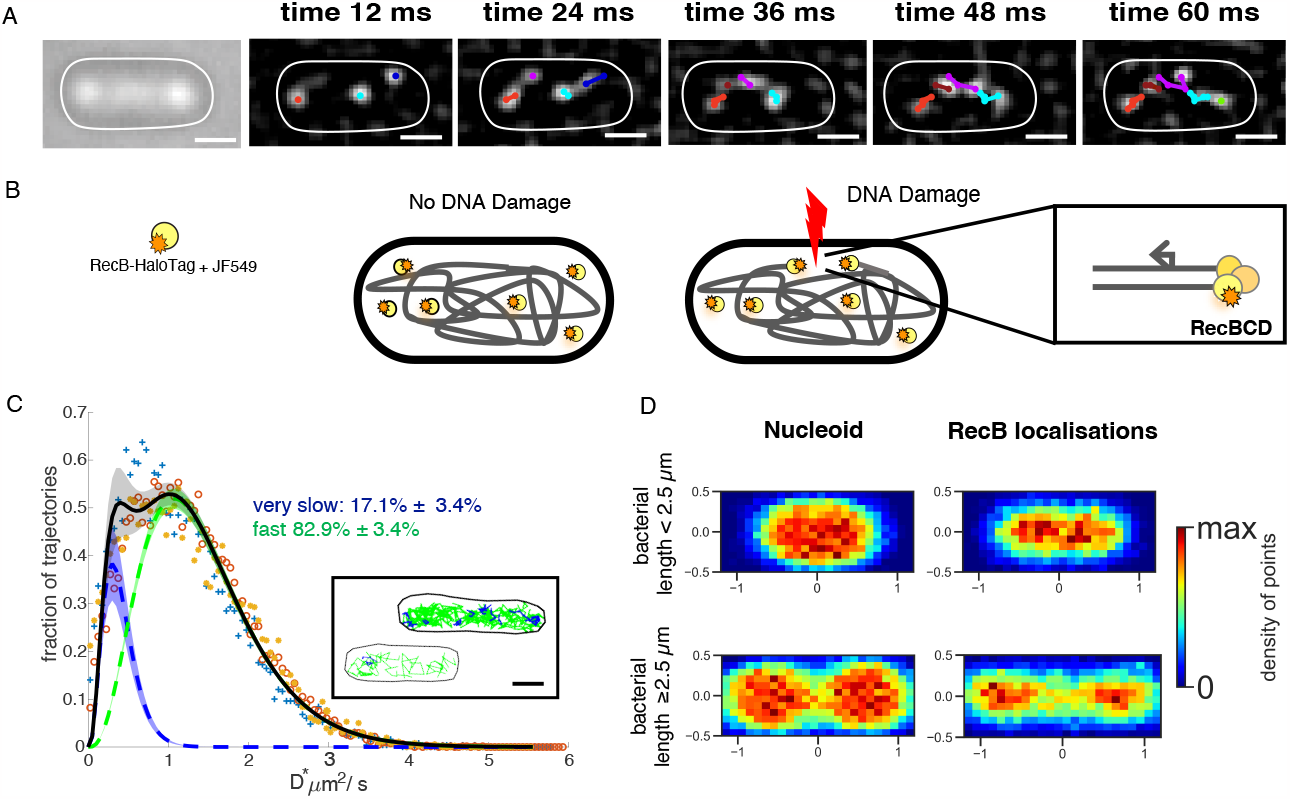
Single RecB molecule tracking in live bacteria. **(A)** Illustration of RecB single-molecule trajectories detected in a single bacterial cell for 5 consecutive frames. Left panel: bright-field. Right panels: progression of the track building overlapped on filtered images to highlight diffraction-limited spots of the frame corresponding to the indicated time (top of each panel). Scale bar 1 µm. **(B)** Schematic of RecB-HaloTag mobility labeled with JF549. In the absence of DNA damage (on the left), RecB-HaloTag mainly undergoes free diffusion. Following DNA damage (on the right), RecB-HaloTag binds to DSB ends. **(C)** Apparent diffusion coefficients distribution, D*, of the detected RecB single-molecule trajectories for three datasets (Total number of bacteria: 2830; Total number of tracks: 25134; dataset 1: ‘+’(blue), dataset 2:’o’(red), dataset 3:’*⋆*’(orange), see Supp. Table 2 for information on single datasets). The averaged fitted distribution describing two sub-populations of RecB trajectories with different mobility is overlapped (full black line). Dotted lines represent the averaged fitted curves, while shaded areas denote the standard deviation from the average of the fits conducted on each dataset. Fractions of trajectories described by each sub-population are indicated. Inset: representative examples of RecB detected trajectories colour-coded as the respective D* sub-population. In blue, are the trajectories whose D* is associated with the slower sub-population, and in green are the ones whose D* is associated with the faster sub-population. Scale bar 1µm **(D)** Localization maps of bacterial cells and nucleoids for cell length smaller (top panels) and equal or longer than 2.5 µm (bottom panels), each normalized by bacterial cell length. Nucleoids N_*cells*_=124; RecB localizations N_*cells*_=1127.

#### 2.6.1 Fluorescent signal acquisition

Images were acquired on the same microscope set-up described above. The GFP signal was excited with a SpectraX Line engine (Lumencor) and a Fluorescein Isothiocyanate Filter (FITC). The exposure time was 200 ms and the camera EM gain was 4. To identify cells within the region of interest (ROI), we obtained 7 bright-field z-stack images around the focal point, covering a total distance of 1.5 *µ*m, with each step being 0.2 *µ*m.

**Table 2:** Percentage of bacteria with RecB on the DNA and high SOS induction for the *recB-HaloTag* and *recB1080-HaloTag* strains.

| Strain | Ciprofloxacin<br>(ng/ml) | RecB on the<br>DNA substrate<br>(% of bacteria) | High SOS<br>induction (% of<br>bacteria) |
| --- | --- | --- | --- |
| *recB-HaloTag* | 0 | 12.3 ± 2.8 | 2.8 ± 0.4 |
| | 4 | 28.8 ± 9.0. | 59.5 ± 13.8 |
| *recB1080-HaloTag* | 0 | 46.6 ± 12.4 | 38.6 ± 10.7 |
| | 4 | 67.5 ± 14.2 | 50.8 ± 3.2 |

#### 2.6.2 Image analysis

Bright-field z-stack and fluorescent images were analysed using the pipeline presented in [32]. In brief, bacterial cells in the ROI were segmented using an edge detection algorithm combined with a custom low-pass filter. The resulting cell outlines underwent manual curation to finalize the segmentation. The fluorescent signal was quantified within each cell, and the average fluorescent signal was calculated by averaging the total signal over the cell area. A local background was computed and subtracted from the average fluorescence, with the background being determined as the average fluorescent signal measured over an area located 15 pixels away from the cell border. Only bacterial cells that were at least 15 pixels away from other cells were included in the analysis, and other cells were excluded.

### 2.7 Single-Particle Tracking

Single-particle tracking was performed using custom-written MATLAB software (MathWorks R2021a^®^). Single-particle localizations were identified by applying an intensity threshold and a bandpass filter to each frame of the video. The coordinates of each intensity peak centroid were computed using a Gaussian fit. Bacterial cell segmentation was used to associate the computed localizations with individual bacterial cells. Trajectories were built inside each segmented bacterium. Localizations within a tracking window of 5 pixels (0.53 *µ*m) in successive frames were linked together to form a trajectory. In the case of multiple localizations in the tracking window, positions whose distance resulted in the minimal total squared displacement were associated with the same track.

The bandpass filter, peak-finder, and tracking functions are from previously developed and published software (http://physics.georgetown.edu/matlab/) [39].

### 2.8 Apparent Diffusion Coefficient Calculation

The D* was calculated as in [40, 41] from the Mean Square Displacement (MSD) of each trajectory divided by four times the time interval between frames, as:

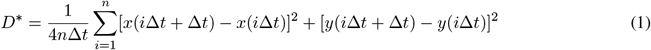

where *x*(*t*) and *y*(*t*) are the trajectory’s position coordinates at time t, the camera exposure time is Δ*t* and *n* is the number of frames. Trajectories were truncated at a fixed length of *n* = 4 frames (5 localizations) to allow the comparison of D* values and to fit an analytical expression to describe the distribution of D*. The localization error was also taken into account and subtracted from the D* values [40].

#### 2.8.1 Localization error

The average localization uncertainty in our experimental conditions was estimated using the Thunderstorm plugin in Fiji[42] on three representative single-particle tracking datasets. The formula used for localization uncertainty is:

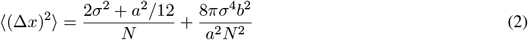

with *σ* the standard deviation of the fitted Gaussian in nm, *a* the pixel size in nm, N the number of detected photons, and *b* the background signal, evaluated as the residuals between the raw data and the fitted Gaussian. The obtained localization uncertainty of 28 nm was used with all datasets to compute the apparent diffusion coefficient.

### 2.9 D* distribution fit

The probability of observing an apparent diffusion coefficient 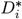for a particle diffusing with *D*^*\**^ and tracked over n frames, is described by the following equation, as established in [43]:

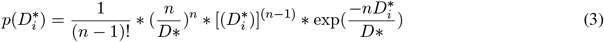

As mentioned above, to compute *D*^*\**^ for each trajectory, we consider trajectories of length *n* = 4 steps (5 localizations). Therefore, to fit the D* histogram distributions, we used the eq.3 for *n* = 4. For the *recB-HaloTag* strain not exposed to ciprofloxacin, we initially fit the *D*^*\**^ with a model describing two molecular species diffusing with 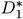 and 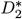:

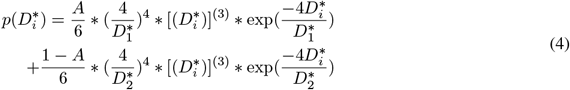

To describe the *D*^*\**^ distribution after exposure to the antibiotic, we used a model describing three different sub-populations of molecules:

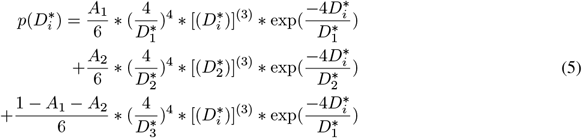

The three-sub-population fit was performed by constraining the value of the averaged *D*^*\**^ of the very slow fraction of trajectories to 0.09 µm^2^/s. This specific value was determined by averaging the *D*^*\**^ values associated with the slower sub-population, which were computed by fitting the *D*^*\**^ distributions of the 14 ng/ml samples using a three-sub-population model for *D*^*\**^. Fits were performed using maximum likelihood estimation in MATLAB.

### 2.10 Calculation of the percentage of bacteria with high SOS induction

We assessed the percentage of cells exhibiting high SOS levels by considering those with SOS values exceeding 2 *×* 10^2^ arbitrary units (a.u.). This threshold was chosen based on the sub-population of bacteria expressing high SOS in the *recB-HaloTag* strain that was not exposed to ciprofloxacin (see Supp. Figure 4D).

### 2.11 Data and code availability

The MATLAB code used to perform the data analysis can be found in the *MEKlab Gitlab*. Please contact the corresponding author for a more recent version. The data that support the findings of this study are available upon request.

## 3 RESULTS

### 3.1 In vivo tracking of single RecB molecules using HaloTag labelling

To understand how DNA Double Strand Breaks (DSBs) are recognized and processed by RecBCD, we measured the mobility of single RecB molecules for different levels of DNA damage. We used a previously characterized translational fusion of the HaloTag to RecB, having already established that the labelling is specific and the fusion can be used to image single RecB molecules which are functional [7].

To further test whether the HaloTag fusion influenced the downstream molecular processes of *E. coli* at the single-cell level, we compared the induction of the SOS response in two strains: the wild-type *E. coli* strain and the *E. coli* strain that contained the RecB-HaloTag fusion. Tests conducted at the single-cell level can detect effects that might be neglected when analysing the impact on DNA repair in *E. coli* in whole populations of bacteria [44]. We measured the cell area and evaluated the induction of the SOS response by calculating the mean GFP intensity per bacterial cell using an SOS transcriptional reporter (*PsulA-mGFP*, [32]). This analysis was performed on both the WT *recB* (MEK445) and *recB-HaloTag* strains (MEK707) after incubating both strains with 10 ng/ml of ciprofloxacin for 2 hours. As expected, after the exposure to ciprofloxacin, we observed an increase in the cell area and induction of the SOS response (see Supplementary Figures 1A and 1B). The cell area and the SOS signal distributions of the *recB-HaloTag* strain were similar to those measured for the WT *recB* (see Supplementary Figures 1C and 1D). Thus, our data show that the HaloTag fusion does not affect the capacity of RecBCD to lead to the induction of the SOS response, further confirming that the *RecB-HaloTag* strain can be used to study DNA DSB repair in live *E*.*coli*.

The HaloTag, conjugated with a synthetic dye (here JF549,[45]), enables *in vivo* tracking of rapidly diffusing proteins [46]. Combined with RecB low copy number (on average 4.9 *±* 0.3 [7]), it allowed us to directly track single RecB molecules without needing photoactivation imaging techniques. To estimate the mobility of a single RecB trajectory, we computed its apparent diffusion coefficient, D*, as previously [46, 47, 48, 49, 41].

We first computed the D* distribution of single RecB molecules in cells not exposed to exogenous sources of DNA damage. We initially performed a fit of the D* histograms using an analytical expression of D* [43] representing one diffusing population of molecules (see Supplementary Figure 2). The value of the fitted D* averaged over the values computed for each of the three datasets was 1.22 *±* 0.10 µm^2^/s (Supp. Figure 2). However, we noticed that the one-population fitted distributions shown in Supplementary Figure 2 failed to fully describe the underlying D* histogram, prompting us to use a two-population fit.

The two sub-populations fit identified a first group of RecB trajectory described by an average D*= 0.40 *±* 0.02 µm^2^/s and a second one described by an average D* = 1.43 *±* 0.05 µm^2^/s (as above, the D* values were averaged from fits performed on three datasets acquired in the same conditions, the error is the standard deviation, see Supplementary Figure 2 and Supplementary Table 3). The majority of RecB trajectories (82.9 *±* 3.4 %, see Supplementary Table 3) showed high mobility (D*= 1.43 *±* 0.05 µm^2^/s). This population of RecB trajectories likely corresponds to RecB molecules that are diffusing in the cytoplasm and do not interact with the DNA. The second population of RecB trajectories, described by the average D* = 0.40 *±* 0.02 µm^2^/s corresponded to 17.1 *±* 3.4% of the entire population. As observed in other DNA-interacting proteins under similar imaging conditions [41], the mobility of this fraction of RecB trajectories is too high to be attributed to molecules bound to the DNA. It is thus likely that this subset of RecB trajectories corresponds to a sub-population of molecules engaged in transient interactions with DNA as they search for their target sites. This two-population fit does not take into account very slow RecB trajectories corresponding to RecB bound to DNA in line with the low frequency of endogenous DSBs [4] (see below).

To verify that RecB molecules localized mainly within the bacterial nucleoid, we built a two-dimensional localization map of the detected RecB molecules for all the bacterial cells of our samples (see Figure 1D, right panel). We then used the SYTOX Green dye [35, 41](see also Material and Methods) to label and image the bacterial nucleoid. Comparing the nucleoid positions to the RecB localization distribution map (Figure 1D, left panel), we observed that, as expected, the RecB spatial distribution overlapped with the spatial distribution of the nucleoid. For larger cells where the chromosome has started to segregate prior to cell division (bacterial cells longer than 2.5 *µ*m) and forms a typical bi-lobar shape, the localization of RecB molecules showed a very similar shape indicating that they likely co-localize with the bacterial nucleoid during the cell cycle (see Supplementary Figure 2H-J for more datasets).

### 3.1 RecB mobility decreases with a high level of induced DNA damage

To investigate how different levels of DNA damage could impact the repair process and RecB mobility, we treated bacterial cells with sub-lethal concentrations of ciprofloxacin. We aimed to identify concentrations of ciprofloxacin that would induce the SOS response without leading to cell death. We used spot test assays across a range of ciprofloxacin concentrations from 0 to 20 ng/ml (see Materials and Methods and Supp. Figure 3). We chose 4 ng/ml and 14 ng/ml for the following reasons: at 4 ng/ml, low-level DNA damage is produced but viability is not affected (see Supp. Figure 3); at 14 ng/ml, the level of DNA damage is higher but the spot tests showed a limited reduction in cell viability (see Supp. Figure 3), thus allowing us to observe the repair process. We quantified the average GFP signal per bacterial cell from the SOS fluorescent reporter *PsulA-mGFP* at the single cell level and measured the cell area in these conditions. Bacteria were exposed to each concentration of ciprofloxacin for 150 minutes before starting microscopy and the same antibiotic concentration was maintained on the agar pad, thus ensuring that the cells reached a “steady-state” of DNA damage exposure. We verified that the two concentrations we selected together with the control represented three distinct levels of SOS induction: no induction except for a few cells (corresponding to endogenous damage) without ciprofloxacin and a low and high level respectively for 4 and 14 ng/ml (Supp. Figure 4D).

We performed RecB single-molecule tracking under ciprofloxacin exposure. As expected, following ciprofloxacin treatment, the bacterial cells appeared elongated (Supp. Figures 4A, 4C), and we observed that the detected RecB trajectories (Figure 2A) explored a smaller space compared to the condition without ciprofloxacin (Figure 1A). This suggests that some RecB molecules were recruited to the DNA and hence appeared much less mobile.

**Figure 2.**
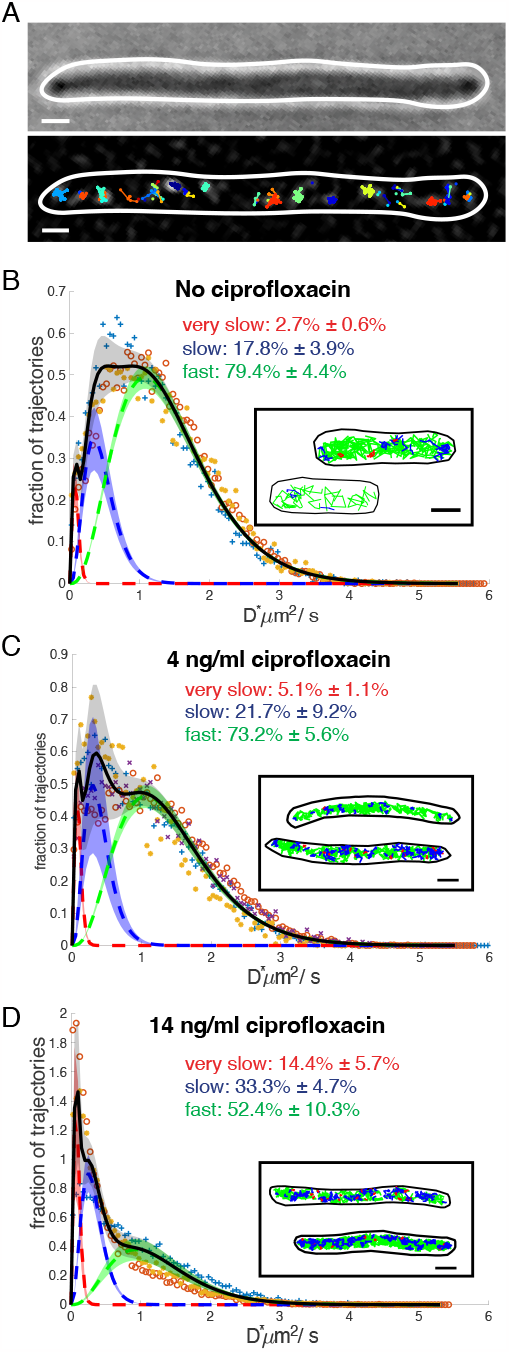
RecB mobility decreases with a high level of DNA damage. **(A)** Representative example of RecB trajectories detected in a cell exposed to 14 ng/ml of ciprofloxacin. Top panel: bright-field. Bottom panel: detected trajectories overlapped on one filtered image. Scale bar 1 µm. **(B)** Apparent diffusion coefficient distributions, D*, for samples exposed to 0 (same datasets shown in Figure 1 here fitted with a three sub-populations model, dataset 1: ‘+’ (blue), dataset 2:’o’(red), dataset 3:’*⋆*’(orange)), **(C)** 4 (dataset 1: ‘+’(blue), dataset 2:’o’(red), dataset 3:’*⋆*’(orange), dataset 4: ‘x’ (purple),) and **(D)** 14 ng/ml (dataset 1: ‘+’ (blue), dataset 2:’o’ (red), dataset 3:’*⋆*’ (orange)) of ciprofloxacin. Histograms were fitted with a three-species model (full black line) corresponding to three sub-populations of very slow (dotted red line), slow (dotted blue line) and fast (dotted green line) moving RecB molecules. Dotted lines are averaged fitted values and shadow areas represent the standard deviation computed from the datasets acquired in each condition. Fractions of trajectories described by each sup-population are indicated. Insets: representative examples of RecB detected trajectories for the corresponding condition, colour-coded as the respective D* sub-population. In red the trajectories described by D*_*veryslow*_; in blue the trajectories described by D*_*slow*_, in green the trajectories described by D*_*fast*_. Scale bar 1µm

The D* distributions for 4 and 14 ng/ml (Figures 2C and D) showed a clear shift toward values of D* smaller than 1 µm^2^/s in comparison to the D* distribution computed for the no ciprofloxacin sample. We also noticed a peak (more evident in the 14 ng/ml of ciprofloxacin condition) in the D* distribution for D* values lower than *∼*0.10 µm^2^/s (Figures 2D and 2C). Similar values of D* have been previously associated with molecules bound to the DNA[40, 48, 50, 41]. We therefore fitted the D* histograms of trajectories obtained in the presence of 14 ng/ml ciprofloxacin with an analytical expression of D* containing a third, additional sub-population of RecB trajectories (See Figure 2, Supp. Figure 5 and Supp. Table 4). The fit was performed by constraining the value of the averaged D* of the very slow fraction of trajectories to 0.09 µm^2^/s (see Materials and Methods). The fit provided an estimate for the relative proportions of each sub-population of RecB trajectories: 14.4 *±* 5.7 % of the detected trajectories were “very slow”; 33.3 *±* 4.7 % were “slow” (D*_*slow*_= 0.33 *±* 0.02 µm^2^/s) and the remaining 52.4 *±* 10.3 % were “fast” (D*_*fast*_= 1.24 *±* 0.07 µm^2^/s). To quantify how the relative fraction of RecB trajectories in the sub-populations changed for the different levels of DNA damage, we performed the same fit for the D* distribution computed for the bacteria exposed to no and 4 ng/ml of ciprofloxacin (See Supp. Table 4 and Supp. Figure 5 for single datasets). We observed that the fraction of trajectories corresponding to D*_*veryslow*_ progressively increased with the level of DNA damage from 2.7 *±* 0.6 % without ciprofloxacin to 5.1 *±* 1.1 % at 4 ng/ml of ciprofloxacin to reach 14.4 *±* 5.7 % at 14 ng/ml of ciprofloxacin. The fraction of RecB trajectories corresponding to the fast sub-population did not vary significantly between 0 and 4 ng/ml of ciprofloxacin with values between 79.4 *±* 4.3 % and 73.2 *±* 5.6 % respectively, but it decreased for the highest level of DNA damage to 52.4 *±* 10.3% (see Supp. Figure 5 and Supp. Table 3). The reduction in the fraction of fast-non-DNA interacting molecules for the higher concentration of ciprofloxacin was consistent with an increase of RecB molecules engaged in the repair process. Taken together, these results suggest RecB mobility experiences small variations at lower concentrations of ciprofloxacin, whereas it is significantly affected at a high ciprofloxacin concentration. This reflects the expected increase in the number of DSBs correlated with increasing ciprofloxacin concentrations.

### 3.3 RecB nuclease inactivation affects DSB repair

To investigate how DSBs are processed when the RecBCD-dependent repair pathway is affected, we chose to observe RecB activity and SOS induction in the presence of a mutated RecB protein, RecB1080. This protein carries a single point mutation in the putative Mg^2+^ binding site of the RecB subunit (Asp-1080 *→* Ala), which inactivates the RecB nuclease domain [21, 22]. Biochemical analysis of RecB1080CD shows that the complex still recognises *χ* sites but does not promote RecA loading [22]. Hence, it is not able to complete DSB repair through the usual RecBCD-dependent RecA loading pathway although it is still partially functional since the helicase activities are not affected. To study this mutant at the single-molecule level, we constructed a HaloTag fusion to the mutated RecB subunit and introduced the SOS transcriptional reporter *PsulA-mGFP* into the mutant chromosome (MEK2324), as previously performed for the *recB-HaloTag* strain. Characterization of this strain (referred to as the *recB1080-HaloTag* mutant) showed normal viability (Supp. Figure 6A and Supp. Figure 8). Survival after exposure to various concentrations of ciprofloxacin (4, 14 ng/ml, see Supp. Figure 6A), whilst reduced compared to WT, was much higher than in a Δ*recB* strain. This is similar to previous results obtained after exposure to gamma irradiation [23] and suggests that the *recB1080-HaloTag* mutant is able to repair DSBs by loading RecA through another, less efficient, pathway (described in Figure 3, [23]). Hence, this mutant allowed us to measure the efficiency of DNA repair when the main RecA loading pathway is inactivated without confounding factors linked to the low viability of most *recBCD* mutants.

**Figure 3.**
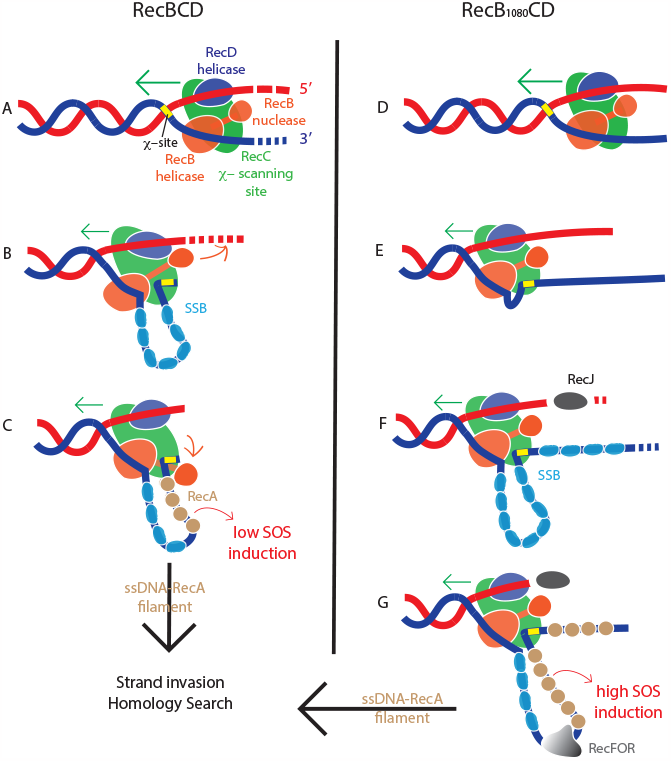
Schematics of the repair pathways leading to RecA loading on ssDNA, based on the models presented in [15, 18, 23]. **(A)** RecBCD (left) translocates on DNA while degrading it. **(B)** After *χ*-site recognition, RecBCD changes its biochemical activity. It pauses and then restarts translocation at a reduced rate; since the *χ* sequence is bound to the RecC subunit the 3’ end exit is blocked. As a result, a loop of ssDNA is accumulated upstream of the *χ*-site and it is rapidly covered by the SSB protein. The 5’ is cleaved more rapidly by the RecB nuclease. **(C)** The RecB nuclease domain promotes the recruitment of the RecA protein to the ssDNA, leading to RecA filament formation. Then, the RecA-ssDNA performs the homology search. **(D)** RecB1080CD (right) translocates on the dsDNA but does not degrade it; **(E)** RecB1080CD recognizes the *χ*-site but does not promote RecA loading. It pauses and possibly undergoes translocation at a reduced rate. Similarly to RecBCD, a loop starts to form upstream of the *χ*-site. **(F)** The 3’ ssDNA is covered by SSB. The 5’ end is degraded by RecJ. Other exonucleases, such as ExoVI [51], could partially degrade the 3’ ssDNA. **(G)** Since RecB1080 lacks nuclease activity, it cannot promote RecA loading, and its continued translocation on the ssDNA could result in a longer 3’ end ssDNA. RecA loading is facilitated by RecFOR. After the RecA-ssDNA filament is formed, it performs strand invasion and homology search.

We first checked the induction of the SOS response in the *recB1080-HaloTag* mutant, using our SOS fluorescent reporter *PsulA-mGFP* and compared it to the *recB-HaloTag* strain. Interestingly, without ciprofloxacin, the SOS signal distribution in the *recB1080-HaloTag* mutant was very different than in the *recB-HaloTag* strain (Figure 4A and Supp. Figures 4A, 4B, 7A and 7B): in the mutant, a large number of cells had induced a detectable level of SOS. Given that both strains are expected to incur the same amount of DSBs (as RecB has no role in DSB formation), this result suggests that WT RecB repairs most endogenous damage very efficiently without inducing the SOS response but that the alternative pathway for RecA loading used in the *recB1080-HaloTag* strain leads to less efficient repair and a higher number of cells inducing SOS as a result. After 150 minutes of exposure to a ciprofloxacin concentration of 4 ng/ml, both the *recB1080-HaloTag* and the *recB-HaloTag* strains had reached approximately the same level of SOS expression suggesting that whilst the repair pathway is affected in the *recB1080-HaloTag* mutant, this strain is still able to induce the DNA damage response upon exposure to exogenous damage.

**Figure 4.**
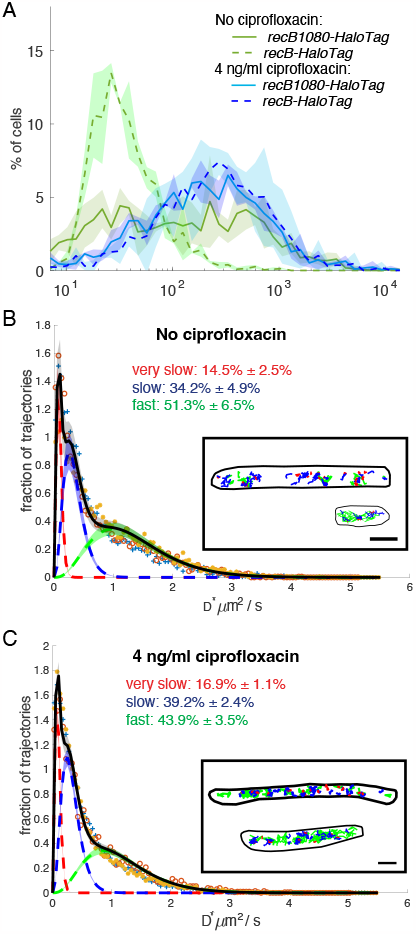
The DNA repair dynamics is altered in *recB1080-HaloTag* mutant.**(A)** In *recB1080-HaloTag* a large fraction of cells induce SOS. Full lines represent the average computed over three datasets, shaded areas represent the standard deviation;**(B)** Apparent diffusion coefficient distributions, D*, for samples treated with no ciprofloxacin (Total number of bacteria: 1232; Total number of tracks: 16444; dataset 1: ‘+’(blue), dataset 2:‘o’(red), dataset 3:‘*⋆*’(orange), see Supp. Table 5 for information on single datasets) and **(C)** 4 ng/ml of ciprofloxacin (Total number of bacteria: 1132; Total number of tracks: 26861; dataset 1: ‘+’(blue), dataset 2:’o’(red), dataset 3:‘*⋆*’ (orange), see Supp. Table 5 for information on single datasets). The averaged fitted distribution describing three sub-populations of RecB trajectories with different mobility is overlapped (full black line). Dotted lines are averaged fitted values and shadow areas represent the standard deviation computed from the datasets acquired in each condition. Fractions of trajectories described by each sup-population are indicated. Insets: representative examples of RecB1080 detected trajectories for the corresponding condition, colour-coded as the respective D* sub-population. In red, trajectories corresponding to D*_*veryslow*_; in blue trajectories corresponding to D*_*slow*_, in green, trajectories corresponding to D*_*fast*_. Scale bar 1µm

To further characterize the *in vivo* activity of RecB1080-HaloTag, we performed tracking of single RecB1080-HaloTag molecules. Without ciprofloxacin, the D* distribution of RecB1080-HaloTag was dramatically different from the one observed in the same condition for the RecB-HaloTag strain. Indeed, the fraction of very slow (D*= 0.09 µm^2^/s) RecB1080-HaloTag molecules was 14.5 *±* 2.5 %, 5 - 6 times larger than for RecB-HaloTag with 2.7 *±* 0.6 % (Figures 4B and 2B, and Supp. Tables 6 and 4). The fraction of slow RecB1080-HaloTag molecules (D*= 0.34 *±* 0.2 µm^2^/s) was 34.2 *±* 4.9 % and the third population of fast RecB1080-HaloTag molecules (D*= 1.33 *±* 0.03 µm^2^/s) was 51.3 *±* 6.5 %. Given that the number of DSB per chromosome is not expected to change in the mutant, the higher fraction of very slow RecB1080 molecules suggests that RecB1080 molecules stay bound to DNA ends for a longer time than the WT RecB, resulting in a shift in the proportion of different types of molecule mobility.

After exposure to 4 ng/ml of ciprofloxacin, we observed that the fraction of very slow RecB1080-HaloTag molecules increased to 17 *±* 1 % from 14.5 *±* 2.5 %. The fraction of slow RecB molecules (D*= 0.32 µm^2^/s) was 39.2 *±* 2.4 % and the third population of fast RecB molecules (D*= 1.15 *±* 0.05 µm^2^/s) was 44.0 *±* 3.5 % (Figure 4C). The increase in the proportion of very slow molecules is similar to what is observed in WT-RecB cells at this concentration and is consistent with the low levels of DNA damage caused by this concentration of ciprofloxacin. This suggests that the amount of exogenous DNA damage induced in *recB1080* is likely to be similar to the one induced in the WT.

To further compare the repair pathways in the *recB-HaloTag* and *recB1080-HaloTag* strains, we analysed the data on an individual cell basis. We calculated the proportion of cells with at least one very slow RecB in each strain as well as the proportion of cells that had strongly induced SOS (Table 2). For endogenous DNA damage, in the *recB-HaloTag* strain, approximately 12% of cells showed at least one RecB bound to DNA but only 2.7% of the cells had induced high SOS levels, suggesting that DNA repair is efficient and rarely leads to full SOS induction. In contrast, in the *recB1080-HaloTag* mutant, the proportion of cells with at least one DNA-bound RecB was very high (46.6%) with a similarly high number of cells that had induced SOS. This high proportion of cells with DNA-bound RecB1080 is likely due to a combination of two phenomena: firstly, as shown above, RecB1080 seems to stay bound to DNA for a longer time than WT RecB. Secondly, as an induced SOS state results in a larger cell volume and more DNA per cell, this could lead to a larger number of DSBs per cell. Upon exposure to ciprofloxacin, there was a simultaneous rise in both the proportion of cells with RecB bound to DNA and exhibiting high SOS induction as expected. Taken together, these results suggest that the dynamics of the SOS induction and the repair time scale in the *recB1080-HaloTag* is altered compared to the WT *recB-HaloTag* strain.

## 4 DISCUSSION

### 4.1 The three sub-populations of RecB molecules correspond to different modes of interaction with DNA

In this work, we used single-molecule tracking in live *E. coli* to achieve a quantitative understanding of the initial steps of DSB repair *in vivo*. We observed that RecB mobility patterns change depending on its engagement in the repair process and that more RecB molecules are recruited on DNA as the level of DNA damage increases.

Our data show that RecB binding to DNA correlates with the presence of DNA double-strand ends. When bacteria experience solely endogenous DNA damage, approximately 2.7% of RecB trajectories are very slow, likely corresponding to RecB bound to the DNA. We also observed that approximately 20% of RecB trajectories (see Figure 2B and Supp. Table 4) are slow, suggesting transient interactions with DNA, probably corresponding to RecB molecules engaged in target search, with the rest freely diffusing in the cytoplasm.

Exposure to exogenous DNA damage results in a critical change in the distribution of RecB interactions with its substrate: a significant proportion of RecB molecules display very slow mobility corresponding most likely to DNA-bound molecules, and the fraction of RecB not involved in the repair process decreases as more molecules bind to DSBs. Our observations are compatible with previous studies of the mobility of DNA repair enzymes such as PolI, LigA, and MutS [40, 50]. All these enzymes bind to damaged DNA, and exhibit a comparatively low fraction of molecules bound to DNA in normal conditions and an increase upon exposure to DNA damage. Similarly, recent work investigating RecB dynamics after exposure to mitomycin C showed an increase of molecules with very low mobility after DNA damage (Figure 4 in [30]).

Our observations are based on tracking the RecB subunit, most likely as part of the RecBC or RecBCD complexes which are both known to interact with DNA[52, 15, 3, 53]. Despite their distinct biochemical activities, it is not possible to distinguish them precisely via single-molecule tracking. Their apparent diffusion coefficients, when non-interacting with DNA, are expected to be nearly indistinguishable due to their high molecular weights (MW RecBC: 263 kDa; MW RecBCD: 330 kDa)[54, 41]. Although RecBCD initiates repair from blunt ends much more efficiently than RecBC [55] they both robustly interact with DNA ends [53] making differentiation of the complexes challenging. It is also possible that we may have detected un-complexed individual RecB subunits. We believe these would represent only a small number of the molecules because RecB forms a tight complex with RecC [15].

Moreover, their impact is likely confined to the fast fraction of the trajectories, as RecB alone lacks robust DNA binding activities[56], in contrast to RecBC and RecBCD.

### 4.2 In the *recB1080* mutant, DSB repair dynamics differ from the WT

Combining RecB single-molecule tracking and the measurement of SOS induction in individual cells highlights key differences between the normal repair pathway and the alternative RecA loading mechanisms in the *recB1080* mutant. When WT RecB is produced, the repair of endogenous damage occurs efficiently without triggering a high SOS response. We observe 12% of the cells with a DNA-bound RecB molecule (see Table 2), suggesting that they are undergoing DNA repair, most likely from replication fork collapse [57]. In these conditions, it is likely that a homologous copy of the chromosome is close by, probably enabling efficient repair, with a short ssDNA-RecA filament, which does not lead to a high SOS response (Figure 3C). Indeed, the lifetime of RecA structures has been reported to be proportional to the induced SOS response [28].

In the *recB1080-HaloTag*, nearly 40% of bacterial cells have induced high levels of SOS, approximately 20 times more than observed in the *recBHaloTag* strain (see Table 2) and similar to the proportion of bacterial cells that have at least one DNA bound RecB1080. This suggests that RecB1080 binding to the DNA mainly results in high SOS induction. Indeed, our results indicate that RecB1080 likely remains bound to the DNA for a longer time than RecB. It is possible that the lack of direct interaction with RecA as a result of the mutation in the nuclease domain [18, 19, 20] affects RecB dissociation process. Moreover, as illustrated in Figure 3G, in the absence of its exonuclease activity, RecB1080 fails to degrade the 3’ DNA overhang before *χ* recognition. The longer translocation combined with the lack of DNA degradation would result in a longer 3’ end and a potentially longer ssDNA-RecA filament, leading to high SOS induction.

Surprisingly, when cells are exposed to a ciprofloxacin concentration of 4 ng/ml for 150 minutes, the percentage of bacterial cells with high SOS induction in the *recB1080-HaloTag* mutants is not higher than that in the *recBHaloTag* strain. This could be explained if loading of RecA using the alternative pathway requires more time in the *recB1080-HaloTag* mutant compared to the *recBHaloTag* strain. Consequently, complete SOS induction might not have been achieved within our observation time window. This is consistent with previous population-based measurements of SOS induction [23] which reported slower SOS induction in a *recB1080* mutant.

## 5 CONCLUSIONS

Our observations contribute to the *in vivo* understanding of DNA DSB repair, providing valuable insights into the interaction between the RecBCD complex and DNA, as well as its ability to respond to varying levels of DNA damage. Our data concerning the *recB1080* mutant confirm the hypothesis that an alternative repair pathway is activated when RecB lacks its nuclease activity. By introducing a point mutation in the RecB nuclease domain to inactivate its exonuclease activity, we were able to understand how a relatively limited perturbation in RecBCD activities affects repair efficiency. Our results highlight the importance of the coordinated action of RecBCD helicases and nucleases, along with RecA loading, in achieving rapid and efficient repair. Moreover, such an experimental approach, using minimal, targeted perturbation of a highly coordinated process, could be used to probe other fundamental biological processes *in vivo*.

## 6 Acknowledgements

We wish to thank Joseph Zamith for his contribution to data acquisition. We are deeply grateful to David Leach and Elise Darmon for generously providing the plasmid to construct the *recB1080* strain and for engaging in insightful discussions. We would also like to express our gratitude to Mark Dillingham for sharing his expertise on RecBCD and we are thankful to Hafez El Sayyed for actively participating in the discussion of our results. This work has been supported by Wellcome Trust Investigator Awards (Grant No. 205008/Z/16/Z awarded to M.E.K. and Grant No. 110164/Z/15/Z awarded to A.N.K.), a BBSRC BB/S008012/1 responsive mode award (to A.N.K. and M.E.K.), and a Marie Skłodowska-Curie Personal Fellowship (Grant No. 101063725-BARTAS) awarded to A.L.

## 7 Authors Contributions

M.E.K., A.L. and A.N.K. conceived the experiments. M.E.K., A.N.K., A.L., D.T. and O.J.P. designed the data analysis. A.L., B.A. and L.McL. built the strains. A.L. and D.T. collected and analysed the data. M.E.K, A.L., D.T.,

A.N.K. and O.J.P. discussed the data. A.L. wrote the manuscript’s first draft and created the figures. M.E.K, A.L. and D.T. revised the manuscript. L.McL., as the lab manager, oversaw order placements and maintained the lab organization. M.E.K and A.N.K. provided funding. All authors read, edited and approved the final manuscript.

## References

[1] Friedberg, E. C. (2003) DNA damge and repair. Nature, 421, 436–440.

[2] Wyman, C. and Kanaar, R. (2006) DNA double-strand break repair: All’s well that ends well. Annual Review of Genetics, 40, 363–383.

[3] Dillingham, M. S. and Kowalczykowski, S. C. (2008) RecBCD Enzyme and the Repair of Double-Stranded DNA Breaks. Microbiology and Molecular Biology Reviews, 72(4), 642–671.

[4] Sinha, A. K., Possoz, C., Durand, A., Desfontaines, J. M., Barre, F. X., Leach, D. R., and Michel, B. (2018) Broken replication forks trigger heritable DNA breaks in the terminus of a circular chromosome. PLoS Genetics, 14(3), 1–28.

[5] Kohanski, M. A., Dwyer, D. J., and Collins, J. J. (2010) How antibiotics kill bacteria: from targets to networks. Nature Reviews Microbiology, 8(6), 423–435.

[6] Michel, B. and Leach, D. (August, 2012) Homologous Recombination—Enzymes and Pathways. EcoSal Plus, 5(1).

[7] Lepore, A., Taylor, H., Landgraf, D., Okumus, B., Jaramillo-Riveri, S., McLaren, L., Bakshi, S., Paulsson, J., and Karoui, M. E. (2019) Quantification of very low-abundant proteins in bacteria using the HaloTag and epi-fluorescence microscopy. Scientific Reports,.

[8] Kalita, I., Iosub, I. A., Granneman, S., and Karoui, M. E. (2021) Fine-tuning of RecBCD expression by post-transcriptional regulation is required for optimal DNA repair in Escherichia coli. bioRxiv, p. 2021.10.23.465540.

[9] Kuzminov, A. (1999) Recombinational Repair of DNA Damage in Escherichia coli and Bacteriophage λ. Microbiology and Molecular Biology Reviews, 63(4), 751–813.

[10] Chaudhury, A. M. and Smith, G. R. (1984) Escherichia coli recBC deletion mutants. Journal of Bacteriology, 160(2), 788–791.

[11] Dermić, D., Halupecki, E., Zahradka, D., and Petranović, M. (2005) RecBCD enzyme overproduction impairs DNA repair and homologous recombination in Escherichia coli. Research in Microbiology, 156(3), 304–311.

[12] Baharoglu, Z. and Mazel, D. (2014) SOS, the formidable strategy of bacteria against aggressions. FEMS microbiology reviews, 38(6), 1126–1145.

[13] Kreuzer, K. N. (2013) DNA Damage Responses in Prokaryotes : Replication Forks. Cold Spring Harb Perspect Biol, pp. 1–23.

[14] Bos, J., Zhang, Q., Vyawahare, S., Rogers, E., Rosenberg, S. M., and Austin, R. H. (2015) Emergence of antibiotic resistance from multinucleated bacterial filaments. Proceedings of the National Academy of Sciences of the United States of America, 112(1), 178–183.

[15] Singleton, M. R., Dillingham, M. S., Gaudier, M., Kowalczykowski, S. C., and Wigley, D. B. (2004) Crystal structure of RecBCD enzyme reveals a machine for processing DNA breaks. Nature, 432(7014), 187–193.

[16] Amundsen, S. K., Taylor, A. F., Reddy, M., and Smith, G. R. (2007) Intersubunit signaling in RecBCD enzyme, a complex protein machine regulated by Chi hot spots. Genes and Development, 21(24), 3296–3307.

[17] Smith, G. R. (2012) How RecBCD Enzyme and Chi Promote DNA Break Repair and Recombination: a Molecular Biologist’s View. Microbiology and Molecular Biology Reviews, 76(2), 217–228.

[18] Churchill, J. J. and Kowalczykowski, S. C. (2000) Identification of the RecA protein-loading domain of RecBCD enzyme. Journal of Molecular Biology, 297(3), 537–542.

[19] Spies, M. and Kowalczykowski, S. C. (2006) The RecA binding locus of RecBCD is a general domain for recruitment of DNA strand exchange proteins. Molecular Cell, 21(4), 573–580.

[20] Lucarelli, D., Wang, Y. A., Galkin, V. E., Yu, X., Wigley, D. B., and Egelman, E. H. (2009) The RecB Nuclease Domain Binds to RecA-DNA Filaments: Implications for Filament Loading. Journal of Molecular Biology, 391(2), 269–274.

[21] Yu, M., Souaya, J., and Julin, D. A. (1998) Identification of the nuclease active site in the multifunctional RecBCD enzyme by creation of a chimeric enzyme. Journal of Molecular Biology, 283(4), 797–808.

[22] Anderson, D. G., Churchill, J. J., and Kowalczykowski, S. C. (1999) A single mutation, RecB(D1080A), eliminates RecA protein loading but not Chi recognition by RecBCD enzyme. Journal of Biological Chemistry,.

[23] Ivančić-Baće, I., Peharec, P., Moslavac, S., Škrobot, N., Salaj-Šmic, E., and Brčić-Kostić, K. (2003) RecFOR function is required for DNA repair and recombination in a RecA loading-deficient recB mutant of Escherichia coli. Genetics, 163(2), 485–494.

[24] Taylor, A. F. and Smith, G. R. (1999) Regulation of homologous recombination: Chi inactivates RecBCD enzyme by disassembly of the three subunits. Genes and Development, 13(7), 890–900.

[25] Lesterlin, C., Ball, G., Schermelleh, L., and Sherratt, D. J. (2014) RecA bundles mediate homology pairing between distant sisters during DNA break repair. Nature, 506(7487), 249–253.

[26] Amarh, V., White, M. A., and Leach, D. R. (2018) Dynamics of RecA-mediated repair of replication-dependent DNA breaks. Journal of Cell Biology, 217(7), 2299–2307.

[27] Ghodke, H., Paudel, B. P., Lewis, J. S., Jergic, S., Gopal, K., Romero, Z. J., Wood, E. A., Woodgate, R., Cox, M. M., and Oijen, A. M. (2019) Spatial and temporal organization of reca in the escherichia coli dna-damage response. eLife, 8, 1–37.

[28] Wiktor, J., Gynnå, A. H., Leroy, P., Larsson, J., Coceano, G., Testa, I., and Elf, J. (2021) RecA finds homologous DNA by reduced dimensionality search. Nature, 597(7876), 426–429.

[29] Wiktor, J., Van Der Does, M., Büller, L., Sherratt, D. J., and Dekker, C. (2018) Direct observation of end resection by RecBCD during double-stranded DNA break repair in vivo. Nucleic Acids Research, 46(4), 1821–1833.

[30] Payne-Dwyer, A. L., Syeda, A. H., Shepherd, J. W., Frame, L., and Leake, M. C. (2022) RecA and RecB: Probing complexes of DNA repair proteins with mitomycin C in live Escherichia coli with single-molecule sensitivity. Journal of the Royal Society Interface, 19(193).

[31] St-Pierre, F., Cui, L., Priest, D. G., Endy, D., Dodd, I. B., and Shearwin, K. E. (2013) One-step cloning and chromosomal integration of DNA. ACS Synthetic Biology, 2(9), 537–541.

[32] Jaramillo-Riveri, S., Broughton, J., McVey, A., Pilizota, T., Scott, M., and El Karoui, M. (2022) Growthdependent heterogeneity in the DNA damage response in Escherichia coli. Molecular Systems Biology, 18(5), 1–14.

[33] Merlin, C., McAteer, S., and Masters, M. (2002) Tools for characterization of Escherichia coli genes of unknown function. Journal of Bacteriology, 184(16), 4573–4581.

[34] Wertman, K. F., Wyman, A. R., and Botstein, D. (January, 1986) Host/vector interactions which affect the viability of recombinant phage lambda clones. Gene, 49(2), 253–262.

[35] Bakshi, S., Choi, H., Rangarajan, N., Barns, K. J., Bratton, B. P., and Weisshaar, J. C. (2014) Nonperturbative imaging of nucleoid morphology in live bacterial cells during an antimicrobial peptide attack. Applied and Environmental Microbiology, 80(16), 4977–4986.

[36] Tokunaga M. (2010) Highly inclined thin illumination enables clear single-molecule imaging in cells. Nature Methods, 5(Fall), 1–7.

[37] Ollion, J., Elez, M., and Robert, L. (2019) High-throughput detection and tracking of cells and intracellular spots in mother machine experiments. Nature Protocols, 14(11), 3144–3161.

[38] Ollion, J., Ollion, C., Gassiat, E., Lehéricy, L., and Corff, S. L. (2021) Joint self-supervised blind denoising and noise estimation. arXiv,.

[39] Crocker, J. (1996) Methods of Digital Video Microscopy for Colloidal Studies. Journal of Colloid and Interface Science, 179(1), 298–310.

[40] Uphoff, S., Reyes-Lamothe, R., de Leon, F. G., Sherratt, D. J., and Kapanidis, A. N. (April, 2013) Single-molecule DNA repair in live bacteria. Proceedings of the National Academy of Sciences, 110(20), 8063–8068.

[41] Stracy, M., Schweizer, J., Sherratt, D. J., Kapanidis, A. N., Uphoff, S., and Lesterlin, C. (2021) Transient non-specific DNA binding dominates the target search of bacterial DNA-binding proteins. Molecular Cell, 81(7), 1499–1514.e6.

[42] Ovesný, M., Křížek, P., Borkovec, J., Švindrych, Z., and Hagen, G. M. (2014) ThunderSTORM: A comprehensive ImageJ plug-in for PALM and STORM data analysis and super-resolution imaging. Bioinformatics, 30(16), 2389–2390.

[43] Vrljic, M., Nishimura, S. Y., Brasselet, S., Moerner, W. E., and McConnell, H. M. (2002) Translational diffusion of individual class II MHC membrane proteins in cells. Biophysical Journal, 83(5), 2681–2692.

[44] Landgraf, D., Okumus, B., Chien, P., Baker, T. A., and Paulsson, J. (April, 2012) Segregation of molecules at cell division reveals native protein localization. Nature Methods, 9(5), 480–482.

[45] Grimm, J. B., English, B. P., Choi, H., Muthusamy, A. K., Mehl, B. P., Dong, P., Brown, T. A., Lippincott-Schwartz, J., Liu, Z., Lionnet, T., and Lavis, L. D. (2016) Bright photoactivatable fluorophores for singlemolecule imaging. Nature Methods, 13(12), 985–988.

[46] Banaz, N., Mäkelä, J., and Uphoff, S. (2019) Choosing the right label for single-molecule tracking in live bacteria: Side-by-side comparison of photoactivatable fluorescent protein and Halo tag dyes. Journal of Physics D: Applied Physics, 52(6).

[47] Kapanidis, A. N., Uphoff, S., and Stracy, M. (2018) Understanding Protein Mobility in Bacteria by Tracking Single Molecules. Journal of Molecular Biology, 430(22), 4443–4455.

[48] Stracy, M., Jaciuk, M., Uphoff, S., Kapanidis, A. N., Nowotny, M., Sherratt, D. J., and Zawadzki, P. (2016) Single-molecule imaging of UvrA and UvrB recruitment to DNA lesions in living Escherichia coli. Nature Communications, 7, 1–9.

[49] Stracy, M., Wollman, A. J., Kaja, E., Gapinski, J., Lee, J. E., Leek, V. A., McKie, S. J., Mitchenall, L. A., Maxwell, A., Sherratt, D. J., Leake, M. C., and Zawadzki, P. (2019) Single-molecule imaging of DNA gyrase activity in living Escherichia coli. Nucleic Acids Research, 47(1), 210–220.

[50] Uphoff, S., Lord, N. D., Okumus, B., Potvin-Trottier, L., Sherratt, D. J., and Paulsson, J. (2016) Stochastic activation of a DNA damage response causes cell-to-cell mutation rate variation. Science, 351(6277), 1094–1097.

[51] Ivanković, S. and Dermić, D. (June, 2012) DNA End Resection Controls the Balance between Homologous and Illegitimate Recombination in Escherichia coli. PLoS ONE, 7(6), e39030.

[52] Taylor, A. and Smith, G. R. (1980) Unwinding and rewinding of DNA by the RecBC enzyme. Cell, 22(2), 447–457.

[53] Jason Wong, C., Lucius, A. L., and Lohman, T. M. (September, 2005) Energetics of DNA End Binding by E.coli RecBC and RecBCD Helicases Indicate Loop Formation in the 3-Single-stranded DNA Tail. Journal of Molecular Biology, 352(4), 765–782.

[54] Kumar, M., Mommer, M. S., and Sourjik, V. (February, 2010) Mobility of Cytoplasmic, Membrane, and DNA-Binding Proteins in Escherichia coli. Biophysical Journal, 98(4), 552–559.

[55] Wu, C. G. and Lohman, T. M. (October, 2008) Influence of DNA End Structure on the Mechanism of Initiation of DNA Unwinding by the Escherichia coli RecBCD and RecBC Helicases. Journal of Molecular Biology, 382(2), 312–326.

[56] Boehmer, P. E. and Emmerson, P. T. (1992) The RecB subunit of the Escherichia coli RecBCD enzyme couples ATP hydrolysis to DNA unwinding. Journal of Biological Chemistry, 267(7), 4981–4987.

[57] Michel, B., Sinha, A. K., and Leach, D. R. F. (2018) Replication Fork Breakage and Restart in Escherichia coli. Microbiology and Molecular Biology Reviews, 82(3).

