## Supplementary Information for "*In vivo* single-molecule imaging of RecB reveals efficient repair of DNA damage in Escherichia coli"

[1]Institute of Cell Biology, University of Edinburgh, Edinburgh, UK; [2]Centre for Synthetic and Systems Biology (SynthSys), University of Edinburgh, UK; [3]Laboratory for Optics and Biosciences, École Polytechnique, Institut Polytechnique de Paris, Palaiseau, FR; [4]Laboratory of Genome Integrity, National Cancer Institute (NCI), National Institutes of Health (NIH), Bethesda, MD, USA; [5]Biological Physics Research Group, Kavli Institute for Nanoscience Research, Department of Physics, University of Oxford, Oxford, UK

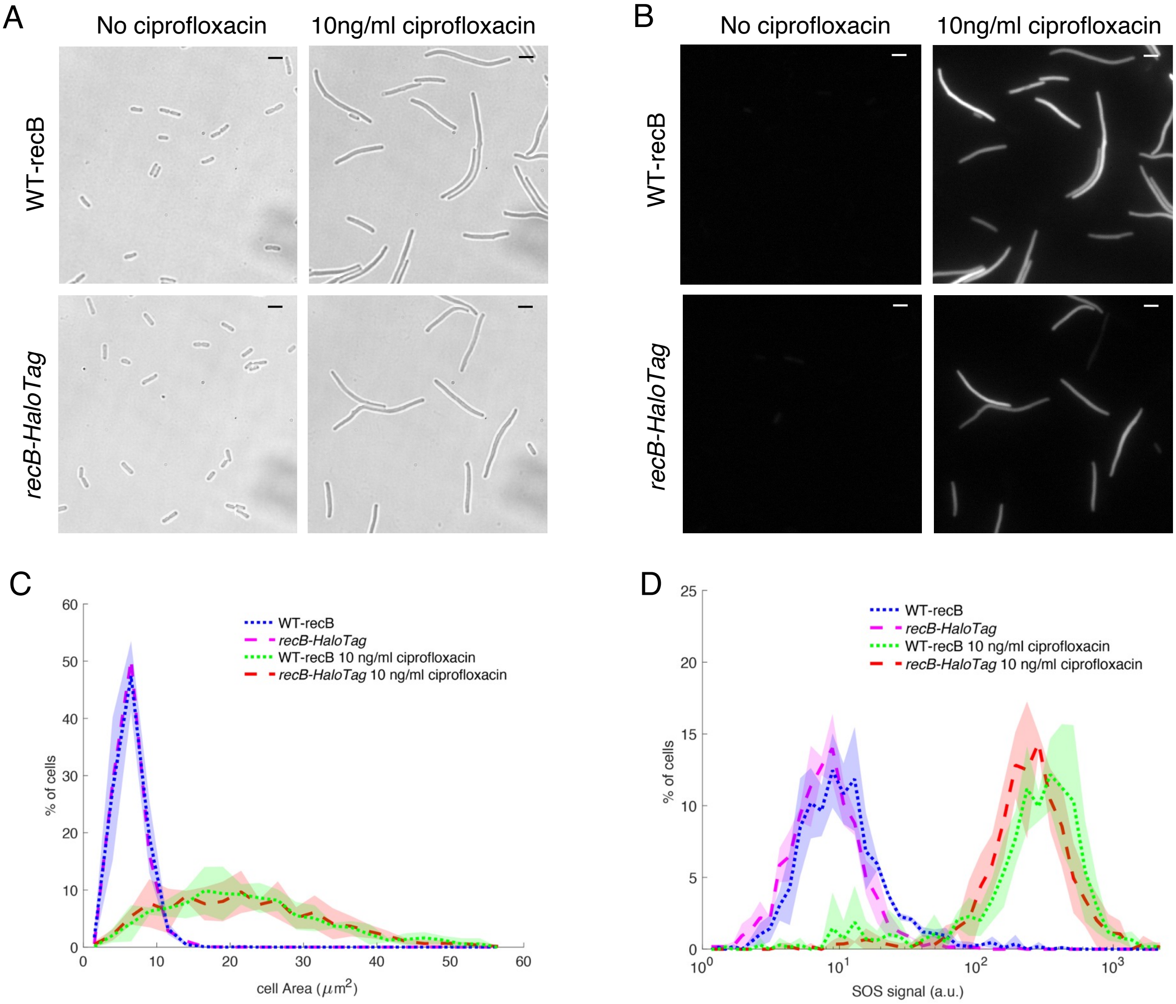


**Supplementary Figure 1: HaloTag fusion to RecB does not perturtb SOS response in bacteria cells** (A) Representative bright-field and (B) SOS induction images of single *E.coli* cells not treated with ciprofloxacin (left top and bottom panels in A and B) and exposed to 10 ng/ml of ciprofloxacin for 150 mins (right top and bottom panels in A and B); top panels: WT-recB strain with SOS reporter *PsulA-mGFP*(MEK455); bottom panels: strain with *recB-HaloTag* fusion and SOS reporter *PsulA-mGFP*(MEK707). Scale bar: 5 μm. (C) Bacterial cells area distributions are the same for WT-recB and *recB-HaloTag* strains in both conditions: sample not exposed to ciprofloxacin and sample treated with 10 ng/ml ciprofloxacin. (D) SOS signal distributions (GFP averaged intensity per bacteria) is the same in both strain for both conditions: sample not exposed to ciprofloxacin and sample treated with 10 ng/ml ciprofloxacin; averages are computed among three technical replica, dotted lines represent the mean of the three dataset, shadow areas represent the standard deviation.

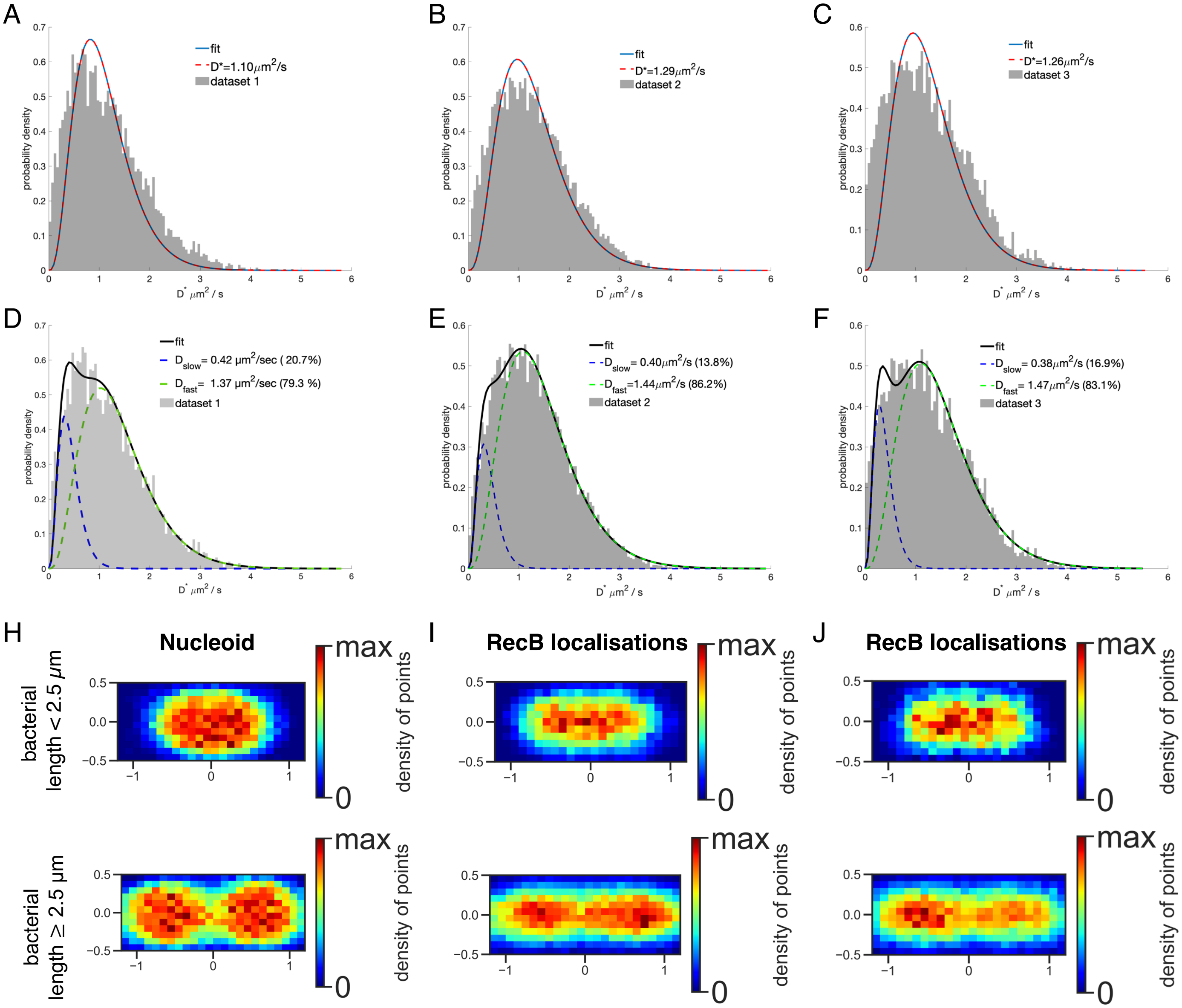


**Supplementary Figure 2: Single datasets of RecB-haloTag D* distribution and localisation maps. (A), (B), (C)** D* histograms of RecB-HaloTag datasets with overlayed fit (red curve) of analytical expression of D* described by one population of RecB molecules. **(D), (E)** and **(F)** D* histograms of same datasets shown in panel **(A), (B)** and **(C)** with overlayed fit (full black curve) of D* described as the sum of two sub-populations of RecB molecules with a $D^*_{slow}$ (blues dotted line) and $D^*_{fast}$ (green dotted line). Number of cells and tracks for each dataset in Supp Table 2; **(H)** Nucleoid localisation distribution of bacterial DNA stained with SYTOX green (Number of bacterial cells: 80) ; **(I)** and **(J)** Localisation maps of bacterial cells of datasets 2 and 3 (same samples as (B), (C), (E) and (F), see Supp. Table 2), top panels (H), (I), (J) : bacterial cells lengths smaller than 2.5 µm ; bottom panels (H), (I), (J) cells lengths equal or longer than 2.5 µm.

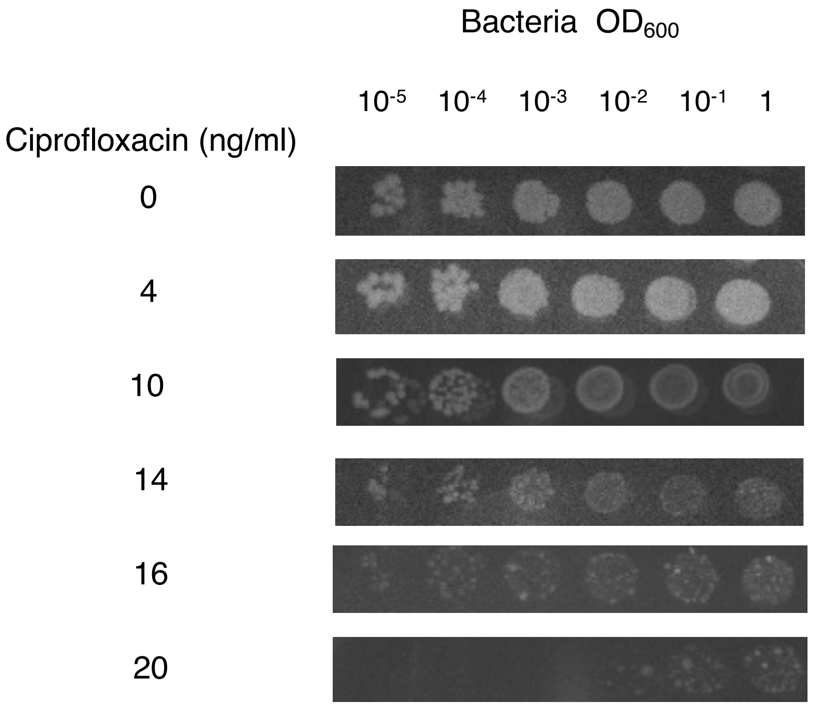


**Supplementary Figure 3: Ciprofloxacin sensitivity test** of *recB-HaloTag* + *PsulA-mGFP* (MEK707) strain for different concentrations of ciprofloxacin (see also Supp. Figure 8).

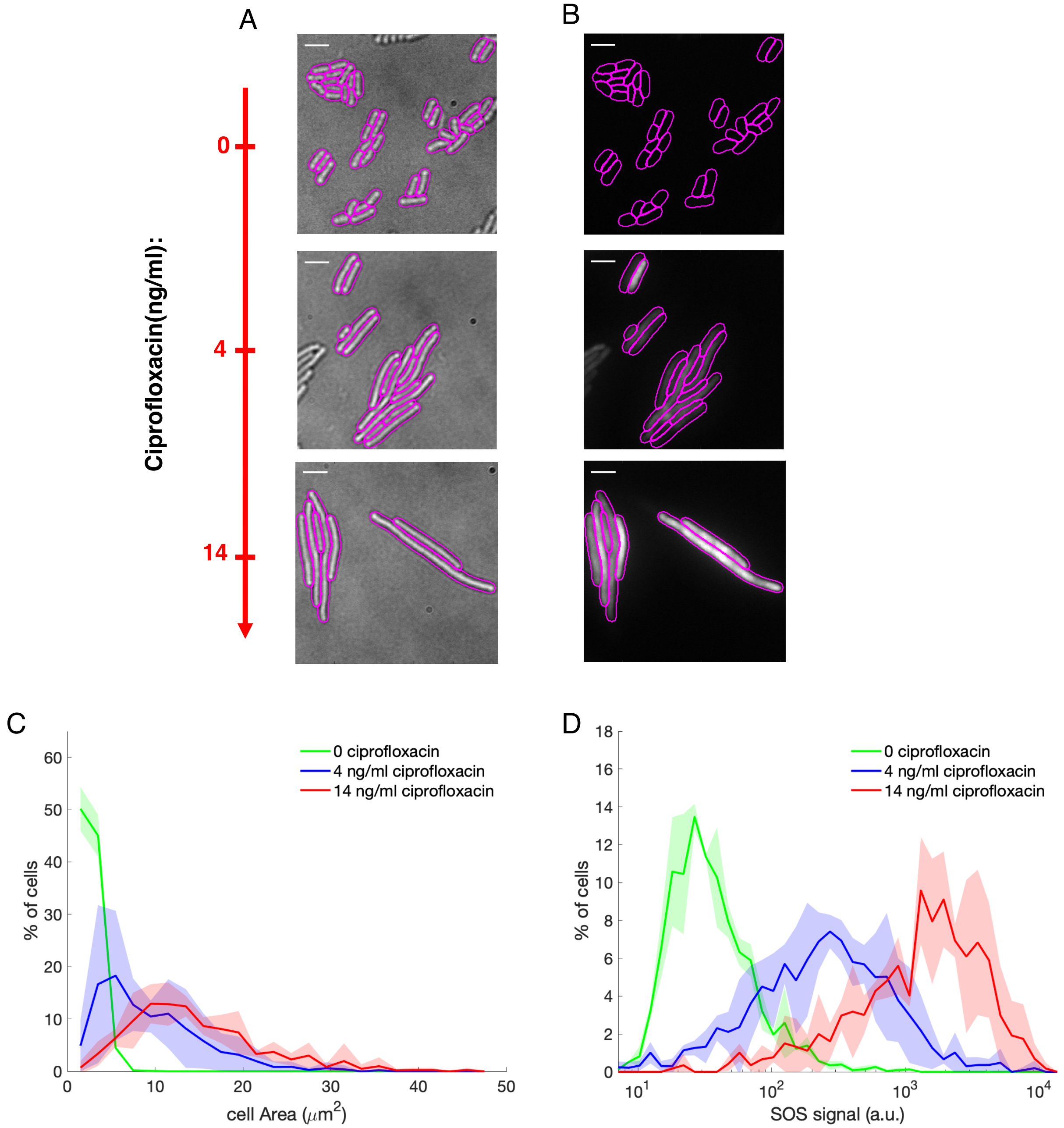


**Supplementary Figure 4**: **Different sub-lethal concentrations of ciprofloxacin induce distinct levels of DNA damage. (A)** Representative bright-field and **(B)** SOS response induction images of single *recB-HaloTag E.coli* cells with SOS reporter *PsulA-mGFP* (MEK707) for increasing concentrations of ciprofloxacin (from top to bottom). Scale bar: 5 µm. **(C)** cell area distributions and **(D)** SOS induction distributions. Averaged datasets: cipro 0: dataset 2 and dataset 3 (dataset 1 has been excluded for an issue in the GFP channel acquisition), cipro 4 and 14 ng/ml: all datasets in Supp. Table 2. Full lines represent the datasets' average, shadow areas the standard deviation.

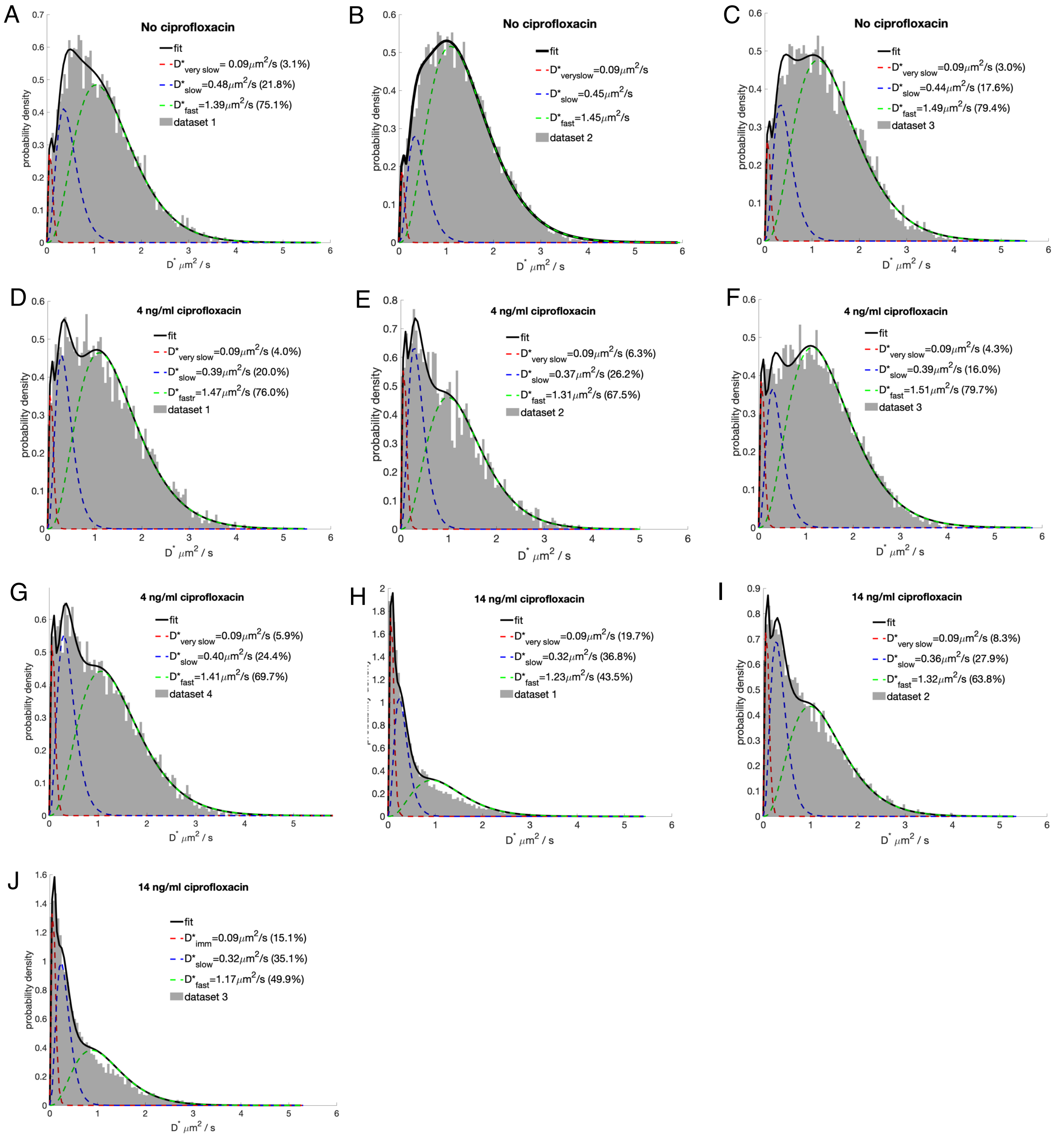


**Supplementary Figure 5: Single datasets of RecB-HaloTag D* distributions at 0, 4, 14 ng/ml ciprofloxacin. (A), (B), (C)** no ciprofloxacin **(D), (E), (G)** and **(F)** 4 ng/ml of ciprofloxacin;**(H), (I), (J)** 14 ng/ml of ciprofloxacin. See also Supp. Tables 2 and 4.

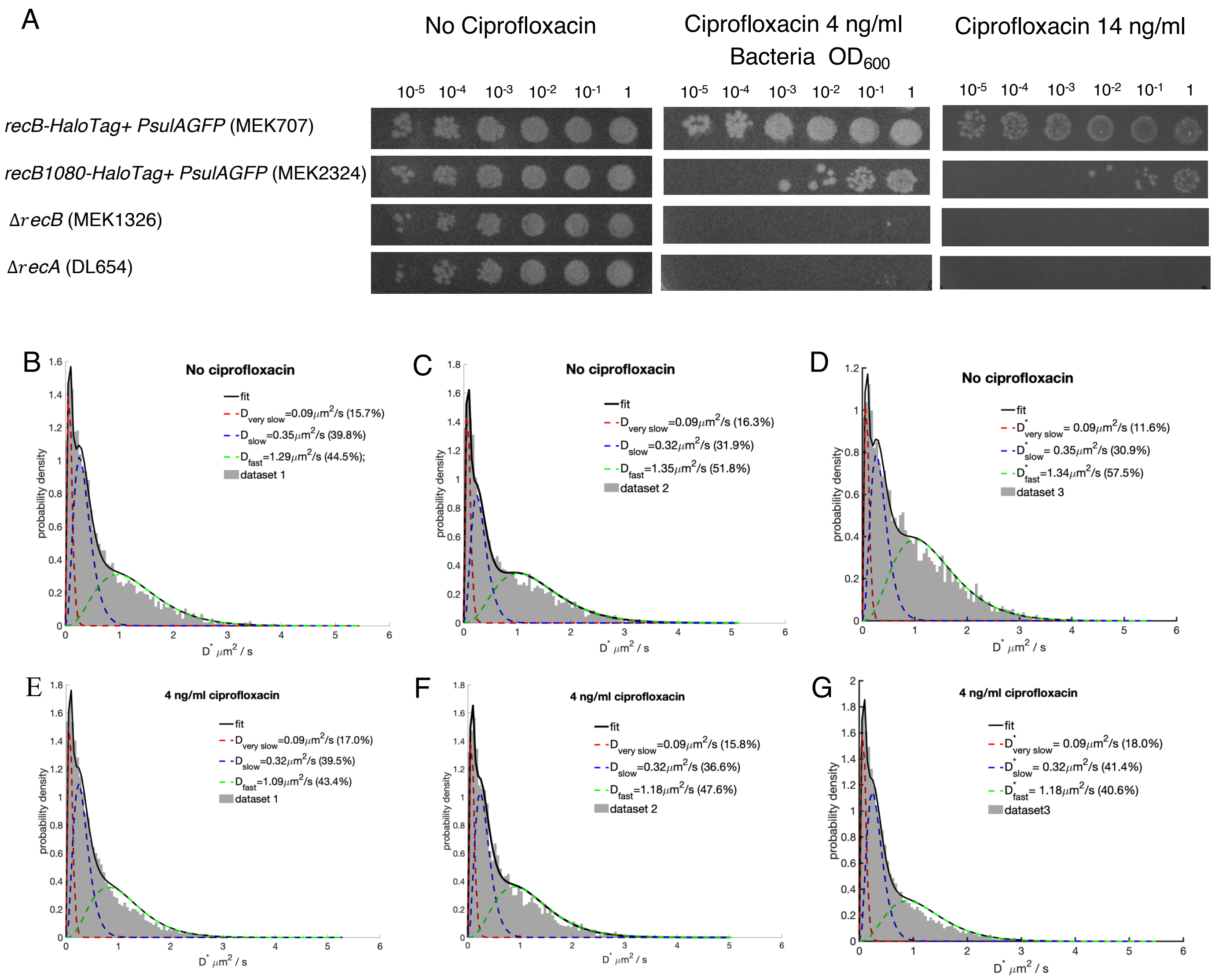


**Supplementary Figure 6**: ***recB1080-HaloTag* mutant. (A) Ciprofloxacin sentitivity tests and single datasets D* distributions at 0, 4, 14 ng/ml ciprofloxacin. (B), (C), (D)** no ciprofloxacin **(E), (F)** and **(G)** 4 ng/ml of ciprofloxacin. (See also Supp. Figure 8 and Supp Tables 5 and 6).

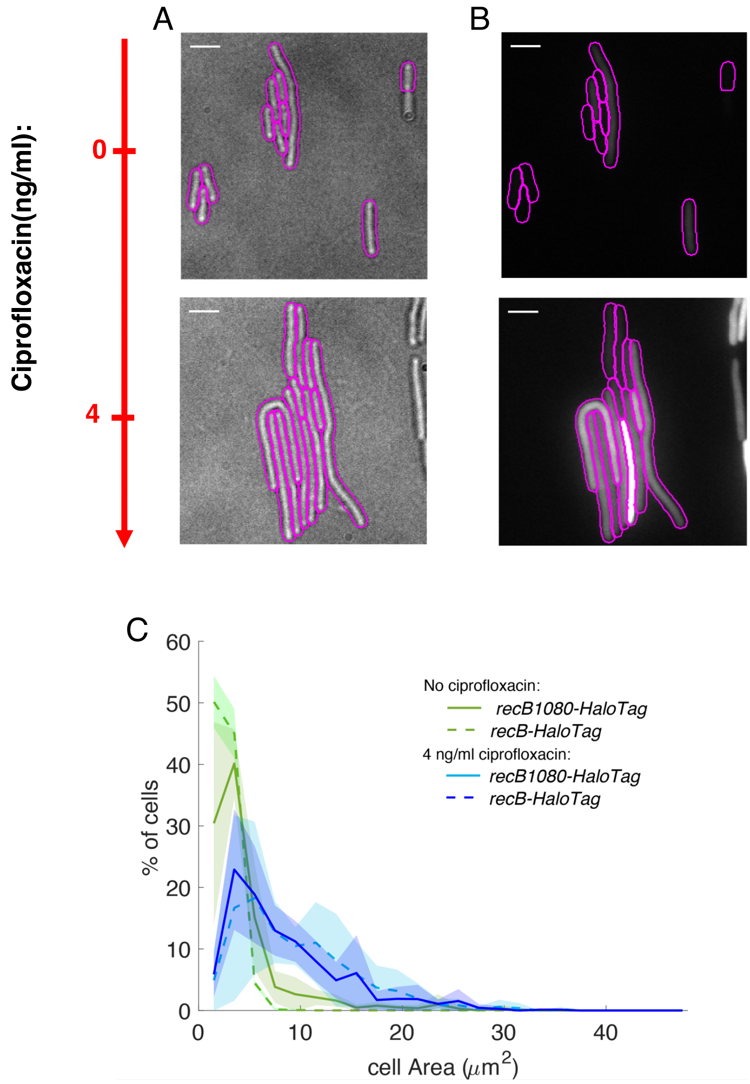


**Supplementary Figure 7**: **RecB1080 SOS induction and cell size (A)** Representative bright-field and **(B)** SOS response induction images of single *recB1080-HaloTag E.coli* cells with SOS reporter *PsulA-mGFP* (MEK2324) exposed to no (top) or 4 ng/mL ciprofloxacin (bottom). Scale bar: 5 µm. **(C)** Cell area distributions. Full lines represent the datasets' average for *recB1080-HaloTag*, dotted lines the datasets' average for *recB-HaloTag*, and shadow areas the standard deviation.

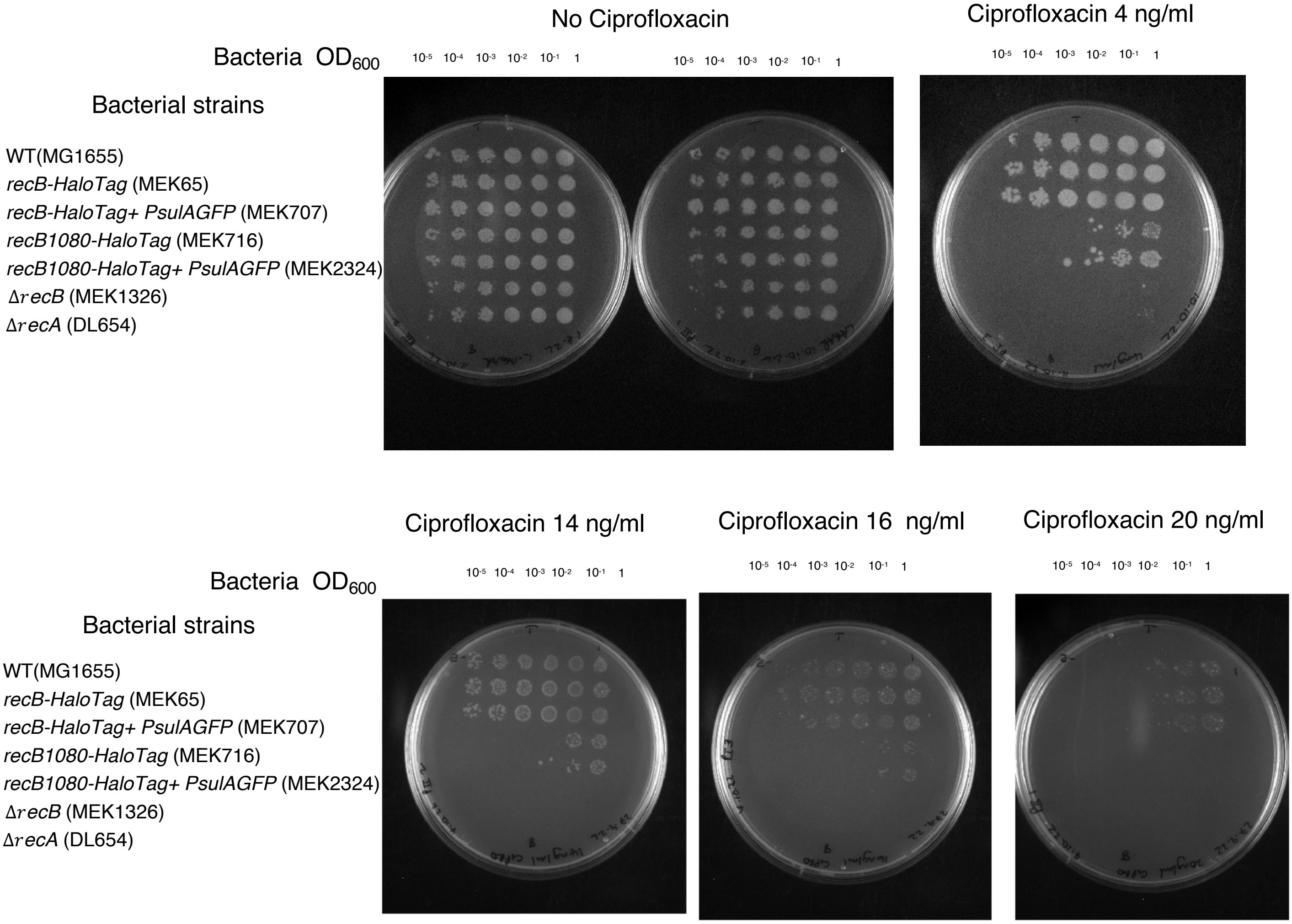


**Supplementary Figure 8**: Ciprofloxacin sensitivity tests (original plates images).

| Primer | 5'- 3' sequence | Purpose |
|---|---|---|
| oSJR058 | GGAATCAATGCCTGAGTG | pOSIP HK022 insertion verification |
| oSJR061 | GGCATCAACAGCACATTC | pOSIP HK022 insertion verification |
| oSJR116 | AGGGTTGATCTTTGTTGT | PsfiA insertion verfication |
| oSJR021 | ATTTAAGAAGGAGATATACAT | mGFP insertion verification |
| Rec1080G-1 | AAAAACTGCAGAACCGACGTTAACACCACATC | pDL4174 construction |
| Rec1080G-2 | GGATTTATAGGCCAGCAGGTA | pDL4174 construction |
| Rec1080G-3 | TACCTGCTGGCCTATAAATCC | pDL4174 construction |
| Rec1080G-4 | AAAAAGTCGACTGTTTGTGCTCCACAGCTTC | pDL4174 construction |
| pKOF | AGGGCAGGGTCGTTAAATAGC | *recB1080* insertion verification |
| pKOR2 | AGGGAAGAAAGCGAAAGGAG | *recB1080* insertion verification |

**Supplementary Table 1** : Oligos used in strain construction

| ciprofloxacin concentration(ng/ml) | Number of bacteria | Number of tracks |
|---|---|---|
| 0 - dataset 1 | 1127 | 5001 |
| 0 - dataset 2 | 1118 | 12937 |
| 0 - dataset 3 | 585 | 7196 |
| 4 - dataset 1 | 288 | 5328 |
| 4 - dataset 2 | 128 | 4036 |
| 4 - dataset 3 | 471 | 11379 |
| 4 - dataset 4 | 305 | 6465 |
| 14 - dataset 1 | 182 | 14874 |
| 14 - dataset 2 | 136 | 10115 |
| 14 - dataset 3 | 192 | 11204 |

**Supplementary Table 2** : *recB-HaloTag* datasets. Number of bacteria and detected tracks for each dataset

| | D* (μm$^2$/sec ) | | % of D* trajectories | |
|---|---|---|---|---|
| ciprofloxacin concentration(ng/ml) | slow | fast | slow | fast |
| 0 - dataset 1 | 0.42 | 1.37 | 20.7 | 79.3 |
| 0 - dataset 2 | 0.40 | 1.44 | 13.8 | 86.2 |
| 0 - dataset 3 | 0.38 | 1.47 | 16.9 | 83.1 |
| 0 - average +/- std | 0.40 +/- 0.02 | 1.43+/- 0.05 | 17.1 +/- 3.4 | 82.9 +/- 3.4 |

**Supplementary Table 3** : two sub-population fit results of RecB-HaloTag D* distributions for all the datasets without ciprofloxacin

| | D* (μm$^2$/sec ) | | | % of D* trajectories | | |
|---|---|---|---|---|---|---|
| ciprofloxacin concentration(ng/ml) | very slow | slow | fast | very slow | slow | fast |
| 0 - dataset 1 | 0.09 | 0.48 | 1.39 | 3.1 | 21.8 | 75.1 |
| 0 - dataset 2 | 0.09 | 0.45 | 1.45 | 2.1 | 14.1 | 83.8 |
| 0 - dataset 3 | 0.09 | 0.44 | 1.49 | 3.0 | 17.6 | 79.4 |
| 0 - average +/- std | 0.09 | 0.46 +/- 0.02 | 1.44 +/- 0.05 | 2.7 +/- 0.6 | 17.8+/- 3.9 | 79.4 +/- 4.4 |
| 4 - dataset 1 | 0.09 | 0.40 | 1.41 | 5.9 | 24.4 | 69.7 |
| 4 - dataset 2 | 0.09 | 0.39 | 1.51 | 4.3 | 16 | 79.7 |
| 4 - dataset 3 | 0.09 | 0.37 | 1.31 | 6.3 | 26.2 | 67.5 |
| 4 - dataset 4 | 0.09 | 0.39 | 1.47 | 4.0 | 20 | 76 |
| 4 - average +/- std | 0.09 | 0.39 +/- 0.01 | 1.42 +/-0.09 | 5.1 +/- 1.1 | 21.7 +/- 9.2 | 73.2 +/- 5.6 |
| 14 - dataset 1 | 0.09 | 0.32 | 1.17 | 15.1 | 35.1 | 49.9 |
| 14 - dataset 2 | 0.09 | 0.36 | 1.32 | 8.3 | 27.9 | 63.8 |
| 14 - dataset 3 | 0.09 | 0.32 | 1.23 | 19.7 | 36.8 | 43.5 |
| 14 - average +/- std | 0.09 | 0.33 +/- 0.02 | 1.24 +/- 0.07 | 14.4 +/- 5.7 | 33.3 +/- 4.7 | 52.4 +/-10.3 |

**Supplementary Table 4** : three sub-population fit results of RecB-HaloTag D* distributions for all the datasets

| ciprofloxacin concentration(ng/ml) | Number of bactiera | Number of tracks |
|---|---|---|
| 0 - dataset 1 | 206 | 5086 |
| 0 - dataset 2 | 326 | 3721 |
| 0 - dataset 3 | 700 | 7637 |
| 4 - dataset 1 | 266 | 12328 |
| 4 - dataset 2 | 329 | 4158 |
| 4 - dataset 3 | 537 | 10375 |

**Supplementary Table 5** : *recB1080-HaloTag* datasets. Number of bacteria and detected tracks for each dataset

| | D* ($\mu m^2$/sec ) | | | % of D* trajectories | | |
|---|---|---|---|---|---|---|
| ciprofloxacin concentration(ng/ml) | very slow | slow | fast | very slow | slow | fast |
| 0 - dataset 1 | 0.09 | 0.35 | 1.29 | 15.7 | 39.8 | 44.5 |
| 0 - dataset 2 | 0.09 | 0.32 | 1.35 | 16.3 | 31.9 | 51.8 |
| 0 - dataset 3 | 0.09 | 0.35 | 1.34 | 11.6 | 30.9 | 57.5 |
| 0 - average +/- std | 0.09 | 0.34 +/- 0.2 | 1.33 +/-0.03 | 14.5+/- 2.5 | 34.2+/- 4.9 | 51.3+/-6.5 |
| 4 – dataset 1 | 0.09 | 0.32 | 1.09 | 17 | 39.5 | 43.4 |
| 4 - dataset 2 | 0.09 | 0.32 | 1.18 | 15.8 | 36.6 | 47.6 |
| 4 - dataset 3 | 0.09 | 0.32 | 1.18 | 18 | 41.4 | 40.6 |
| 4 - average +/- std | 0.09 | 0.32 +/- 0 | 1.15 +/-0.05 | 16.9 +/- 1.1 | 39.2 +/- 2.4 | 43.9+/-3.5 |

**Supplementary Table 6** : three sub-population fit results of RecB1080-HaloTag D* distributions for all the datasets
